# ALS mutations do not alter perineuronal net formation in human stem cell-derived motor neurons

**DOI:** 10.1101/2025.09.13.676004

**Authors:** Caoimhe Kerins, Ivo Lieberam, Eileen Gentleman

## Abstract

Perineuronal nets (PNNs) are extracellular matrix structures that stabilise synaptic inputs and regulate neuronal plasticity. Although PNN dysregulation is observed in several neurological disorders, their relevance to amyotrophic lateral sclerosis (ALS) remains unclear. In particular, the extent to which PNN alterations in ALS are motor neuron (MN)-intrinsic is unknown. We investigated whether human pluripotent stem cell-derived MNs form PNN-like structures *in vitro*, and whether ALS-associated mutations alter this process. We show that MNs generate PNN-like structures containing hyaluronan, tenascin-R, and aggrecan and that their formation and gene expression were not altered by ALS mutations. To explore whether PNN dysregulation reflects contributions from other cell types or selective MN vulnerability, we conducted meta-analyses of transcriptomic datasets from pluripotent stem cell-derived astrocytes carrying ALS-associated mutations, as well as datasets comparing MN populations with differential susceptibility to ALS. These analyses revealed no consistent differences in PNN-related gene expression in human stem cell-derived MN. In contrast, transcriptomic analyses of human post-mortem ALS tissues revealed significant dysregulation of PNN-related genes, including core PNN components and linker proteins. These findings suggest that PNN changes in ALS are not MN-intrinsic, but may result from interactions with other cell types in the central nervous system such as glia.

## Introduction

Amyotrophic lateral sclerosis (ALS) is a degenerative neuromuscular disease characterised by death of motor neurons (MNs) and resultant muscle paralysis. There is currently no cure for ALS and most patients die of neuromuscular respiratory failure within 2-5 years of diagnosis^1^. 80-90% of ALS cases are sporadic and are usually caused by an interplay of environmental and genetic factors^2^. The remaining cases are familial, and predominantly associated with a single dominant mutation in one of ∼30 known ALS-linked genes^3^, including chromosome 9 open reading frame 72 (C9orf72), superoxide dismutase 1 (SOD1), transactive response DNA binding protein 43 (TDP-43) and fused in sarcoma (FUS)^4^. Several genetic causes and risk factors including biological sex and age have been associated with ALS^2^. Moreover, multiple molecular changes have been noted in MNs in ALS models, including oxidative stress, excitotoxicity, neuroinflammation, neuronal hyperexcitation, and axonopathy^1^. Differing explanations have been proposed to explain MN death in ALS. For example, hypotheses commonly labelled “dying forward” and “dying back” propose alternative initiation sites for MN degeneration in ALS^5^. However, both may occur simultaneously or sequentially, and the key mechanism(s) that drive ALS pathology remain elusive. Adding to this complexity, MNs are not uniformly affected in ALS. For example, fast-twitch α-MNs are highly susceptible, whereas other MNs such as those in the oculomotor nucleus remain relatively preserved, even at advanced disease stages^6,7^. These uncertainties underscore the need to consider not only cell-intrinsic mechanisms but also extracellular influences that may shape MN vulnerability in ALS.

Three major extracellular structures make up the extracellular matrix (ECM) of the central nervous system: the basement membrane, perineuronal nets (PNNs), and the diffuse matrix. PNNs directly surround the cell body, proximal dendrites and axon initial segment of some neurons, including many interneurons and MNs^8,9^. PNN components are secreted by neurons, oligodendrocytes, microglia, and astrocytes^10^, and consist of a hyaluronan backbone and multiple proteoglycans including aggrecan, versican, brevican, and other linker proteins, such as tenascin-R. In rat, up to 80% of spinal MNs have PNNs, and 90% of α-MNs are positive for aggrecan^11^. Components of PNNs are liable to degradation by a host of catabolic enzymes including matrix metalloproteinases (MMPs) and members of the disintegrin and metalloproteinase with thrombospondin motifs (ADAMTS) family^12^. Various functions have been associated with PNNs including protection of neurons against oxidative stress, particularly iron-induced oxidative stress^13^, regulation of memory and plasticity^14–16^, and control of neuronal excitability^17,18^. Alterations to PNNs have been reported in a range of neurological diseases such as Alzheimer’s disease, schizophrenia, multiple sclerosis and epilepsy^18–20^.

Several studies have suggested potential links between PNNs and ALS pathology. For example, both TDP-43 Q331K and SOD1 G93A mice show significant reductions in PNNs around α-MNs at disease onset and mid-stage compared to controls^21,22^. Similarly, transgenic rats harbouring a His46Arg mutation in the SOD1 gene, demonstrate a progressive loss of PNNs around susceptible MNs in the ventral horns of the lumbar spinal cord^23^. Additional work suggests that PNN components influence excitatory balance, alterations of which are a pathological hallmark of ALS. For example, depletion of hyaluronan synthase 3 (*Has3*), which is expressed in PNN-bearing MNs in the spinal cord^8^, results in an epileptic phenotype^24^. Excitatory synaptic transmission is also increased in tenascin-R-deficient mice^25^, which also exhibit motor coordination deficits^26,27^. Other studies have implicated matrix remodelling enzymes. For example, whilst MMP9 is expressed by the most vulnerable MNs to cell death in ALS, it is absent in areas that are resistant to disease^28^. Moreover, reducing MMP9 function resulted in SOD1 G93A mice retaining 75% innervation in the normally vulnerable tibialis anterior muscle compared to controls^28^. Evidence in humans is more limited, but post-mortem studies on sporadic ALS patients have identified increases in hyaluronan binding protein 2 and decreases in tenascin-R in their cerebral spinal fluid^29^.

Taken together, these data suggest that maintenance of normal PNNs may help preserve MN function, and that their loss may contribute to ALS pathology. However, testing this hypothesis remains challenging due to key gaps in our understanding of PNNs, particularly in humans. Indeed, there is scant evidence on the extent to which human central nervous system (CNS) cells (such as MNs) form PNNs, if ALS-associated mutations affect PNN production by MNs, and whether PNNs are altered in humans with ALS. Here, we show that human pluripotent stem cell-derived MNs form PNN-like structures *in vitro*, and that ALS mutations do not alter their formation or their expression of PNN-related genes. Similarly, a meta-analysis failed to detect differential expression of PNN-related genes when comparing resistant and vulnerable MN populations or mutant to control human induced pluripotent stem cell (hiPSC)-derived astrocytes. However, a re-analysis of post-mortem transcriptomic data revealed altered expression of PNN components in ALS tissues compared to healthy controls. These findings suggest that PNN disruption in ALS is unlikely to be MN-intrinsic, but may instead involve other CNS cell types such as glia or region-specific factors not captured in *in vitro* models.

## Results

### Human pluripotent stem cell-derived MNs form PNNs *in vitro*

Previous work has shown that primary cultures of rodent neurons and glial cells produce PNN-like structures *in vitro*^30–33^. However, there are currently no studies that report PNN formation by human pluripotent stem cell-derived CNS cells. Therefore, we first established a 2D model consisting of human pluripotent stem cell-derived MNs co-cultured with mouse embryonic stem cells (ESC)-derived astrocytes (Figure 1A). This protocol relies on parallel directed differentiation of human ESC/iPSC to MNs^34^ and mouse ESC to astrocytes^35,36^. To generate co-cultures, we enriched each population using magnetic activated cell sorting (MACS): human MNs were isolated from Hb9::CD14-engineered hESCs/hiPSCs, and mouse astrocytes from GFAP::CD14-expressing ESCs^35–37^. These co-cultures stained positively for β-III tubulin and Glial Fibrillary Acidic Protein (GFAP), confirming the presence of both cell types (Figure 1B).

**Figure 1:**
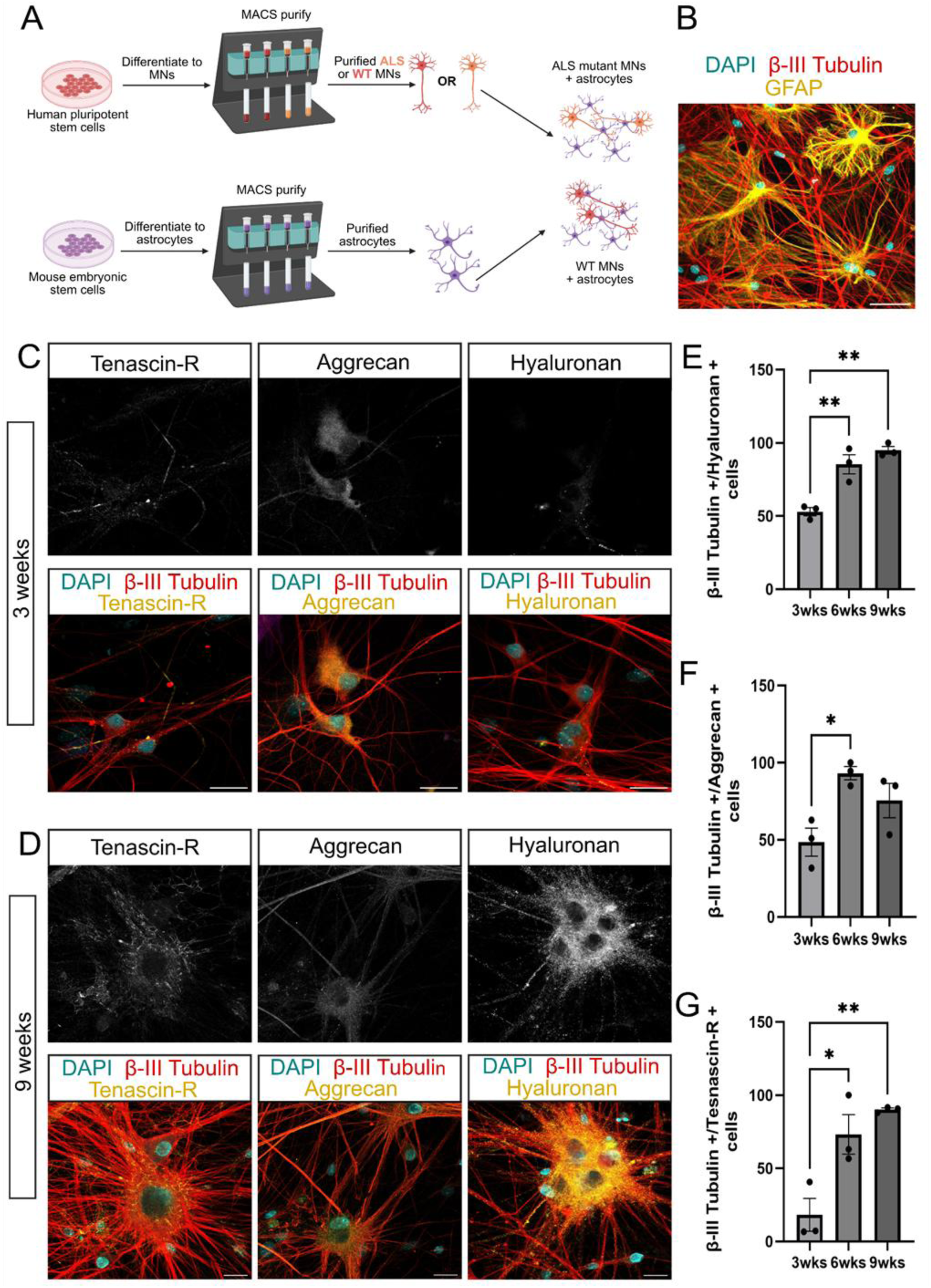
Human pluripotent stem cell-derived MNs form PNNs in *vitro.* PNN-like structures are deposited in co-cultures of WT human ESC-derived MNs and astrocytes. A) Schematic overview of co-culture system. Mouse ESC were differentiated into astrocytes by directed differentiation and human WT ESC (H9) into MNs. At the end of differentiation, MACS was used to purify astrocyte and **MN** populations. Astrocytes were plated 1 week before MNs. Co-cultures were then maintained for either 3, 6 or 9 weeks. B) Representative confocal microscopy image of immunostaining illustrating MNs (β -Ill tubulin) and astrocytes (GFAP) after 3 weeks in co-culture. Representative confocal microscopy images showing C) 3-week and D) 9-week staining of PNN components aggrecan, tenascin-R and hyaluronan in co-cultures of MNs and astrocytes. Scale bar= 25 µm. Quantification of PNN-like structures in co-cultures of MNs and astrocytes at 3, 6 and 9 weeks. Quantification shows the percent of MNs that are double positive for β -Ill tubulin and PNN component E) hyaluronan, F) aggrecan, or G) tenascin-R. N=3 differentiations with 5 images/N. *P<0.05, **P< 0.01; ordinary, one-way ANOVA.

After 3 weeks, immunostaining confirmed that co-cultures were positive for PNN components hyaluronan and aggrecan, which have been shown to be necessary for the formation of a PNN-like matrix *in vitro*^38^; and tenascin-R, which plays essential roles in clustering aggrecan, stabilising nets and promoting the expression of brevican, neurocan and hyaluronan^39^ (Figure 1C). Cultures also stained positively with Wisteria Floribunda Agglutin (WFA) (Figure S1). WFA is a lectin classically used to identify PNNs by binding to an unknown motif in chondroitin sulfate glycosaminoglycan chains. PNN-like structure deposition was not evident at earlier time points (Figure S2). After 9 weeks of co-culture, further accumulation of PNN components was evident (Figure 1D), and there was a striking qualitative difference in the intensity of staining for PNN components in co-cultures after 9 weeks compared to 3 weeks. Cultures continued to stain positively with WFA, which was diffuse in contrast to PNN component aggrecan, which strongly localised with MNs (Figure S3). At 3 weeks, tenascin-R generally deposited around neurites, aggrecan was mostly around cell somata, and hyaluronan appeared punctate and spread around both cell somata and neurites. At later time points, quantification of immunostaining showed increased fractions of hyaluronan, tenascin-R and aggrecan positive cells that were also positive for β-III tubulin (Figure 1E-G, Table S1, and Supplementary Code 1). After 9 weeks, tenascin-R was observed in a linearised/striped pattern around the cell somata of MNs and neurites. Aggrecan was again associated with cell somata as well as neurites while hyaluronan maintained its punctate appearance around MN somata and neurites. PNN-like structures appeared to be mostly MN-derived as they were concentrated around MNs and not astrocytes (Figure S4). Thus, human pluripotent stem cell-derived MNs form PNN-like structures *in vitro* that robustly stain for hyaluronan, aggrecan and tenascin-R.

### ALS mutations do not impact PNN formation by human pluripotent stem cell-derived MNs

Following confirmation that PNN-like structures were formed in *in vitro* co-cultures of MNs and astrocytes, we proceeded to form MNs from TDP-43 G298S and C9orf72 (C9-3) ALS mutant and isogenic control human iPSC lines (see Table 1 for detailed descriptions of the lines) using the same co-culture strategy outlined in Figure 1A. Consistent with our observations in wild-type human ESCs, MNs formed from mutant and control iPSC lines all formed PNN-like structures that stained positively for hyaluronan, aggrecan, and tenascin-R (Figure S5). We then analysed these images to localise MNs (β-III tubulin positive soma and projections) with positive staining for PNN components (aggrecan/hyaluronan/tenascin-R). We found that the fraction of MNs that were double positive for β-III tubulin and either hyaluronan, aggrecan or tenascin-R increased over the 9-week culture (Figures 2A-C and Table S2). However, we could detect no significant differences in the number of double positive MNs between any ALS mutant cell line and its control at any time point (3, 6 or 9 weeks). Therefore, we conclude that TDP-43 G298S and C9-3 ALS mutations do not affect the ability of human pluripotent stem cell-derived MNs to create aggrecan-, tenascin-R- and hyaluronan-containing PNN-like structures *in vitro*.

**Figure 2:**
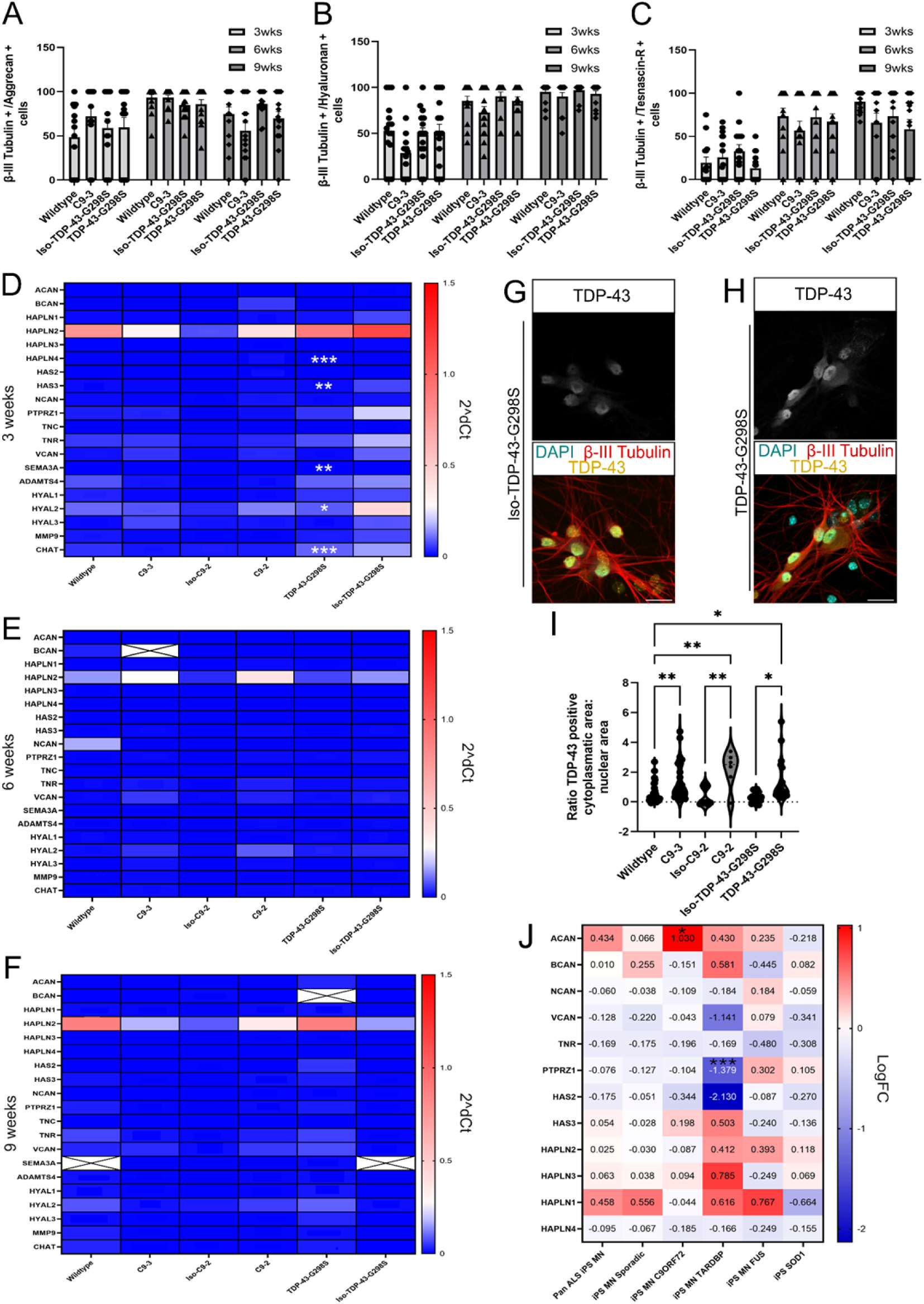
ALS mutations do not impact PNN formation by human pluripotent stem cell derived MNs. Quantification of PNN-like structures in co-cultures of MNs and astrocytes at 3, 6 and 9 weeks. MNs were either WT (H9), C9ORF72 mutant (C9-3), TARDBP mutant (TDP-43) or isogenic control TARDBP (lso-TDP-43). Quantification shows the percent of MNs (β-Ill tubulin+) that were also positively stained for PNN component A) aggrecan, B) hyaluronan, or C) tenascin-R. N=3 differentiations; 5 images/N. Relative gene expression for a range of PNN components and catabolic regulators of PNNs performed using human-specific primers on co-cultures of MNs and astrocytes after D) 3, E)6, and F) 9 weeks in culture. 2^dCt values are relative to housekeeping genes *(GAPDH* or *ACTB)* and significance is indicated when there was a change in gene expression relative to the mutant line’s control (C9-3 compared to WT cultures). Undetected genes are indicated by crossed out boxes. Representative confocal microscopy images of TDP-43 staining in G) TDP-43 G298S isogenic controls and H) TDP-43 G298S mutants in co-cultures. I) Quantification of TDP-43 cytoplasmic/nuclear localisation. *P<0.05, **P< 0.01, ordinary one-way ANOVA. H) Log fold change (LogFC) expression levels of genes encoding various PNN components generated from a meta-analysis comparing MNs derived from iPSC ALS mutant compared to control lines. ***P<0.001.

**Table 1:** List of cell lines and their sources.

| Cell line | Details of source |
| --- | --- |
| H9 human ESC line (Wildtype, WT) | WiCell |
| M211R2 C9orf72-mutant human iPSC line (C9-3) | Mehta *et al. Acta Neuropathol* 2021<sup>46</sup> |
| DN1qV4 C9orf72-mutant human iPSC line (C9-2) | Mehta *et al. Acta Neuropathol* 2021<sup>46</sup> |
| Isogenic DN1qV4 control human iPSC line (Iso-C9-2) | Mehta *et al. Acta Neuropathol* 2021<sup>46</sup> |
| TDP-43 G298S-mutant human iPSC line (TDP-43-G298S) | Barton *et al. Brain Commun* 2021<sup>47</sup> |
| Isogenic TDP-43 G298S control human iPSC line (Iso-TDP-43-G298S) | Harley *et al. Cell Rep* 2023<sup>36</sup> |
| H9 human ESC line, Wildtype, WT, Hb9::CD14 transgenic | Harley *et al. Cell Rep* 2023<sup>36</sup> |
| M211R2 C9orf72-mutant human iPSC line C9-3, Hb9::CD14 transgenic | Harley *et al. Cell Rep* 2023<sup>36</sup> |
| DN1qV4 C9orf72-mutant human iPSC line, C9-2, Hb9::CD14 transgenic | This study |
| Isogenic DN1qV4 control human iPSC line, Iso-C9-2, Hb9::CD14 transgenic | This study |
| TDP-43 G298S-mutant human iPSC line, TDP-43-G298S, Hb9::CD14 transgenic | Harley *et al. Cell Rep* 2023<sup>36</sup> |
| Isogenic TDP-43 G298S control human iPSC line, Iso-TDP-43-G298S, Hb9::CD14 transgenic | Harley *et al. Cell Rep* 2023<sup>36</sup> |
| H9 human ESC line, Wildtype, Hb9::CD14/CAG::CAG::ChR2-YFP transgenic | Harley *et al. Cell Rep* 2023<sup>36</sup> |
| M211R2 C9orf72-mutant human iPSC line C9-3, Hb9::CD14/CAG::CAG::ChR2-YFP transgenic | Harley *et al. Cell Rep* 2023<sup>36</sup> |
| TDP-43 G298S-mutant human iPSC line TDP-43-G298S, Hb9::CD14/CAG::CAG::ChR2-YFP transgenic | Harley *et al. Cell Rep* 2023<sup>36</sup> |
| Isogenic TDP-43 G298S control human iPSC line Iso-TDP-43-G298S, Hb9::CD14/CAG::CAG::ChR2-YFP transgenic | Harley *et al. Cell Rep* 2023<sup>36</sup> |
| IB10 mouse embryonic stem cell (mESC) line, GFAP::CD14/CAG::GDNF transgenic | Bryson *et al., Science* 2014<sup>35</sup> |

Although immunostaining failed to show significant differences in PNN-like structure formation between MNs derived from TDP-43 G298S and C9-3 mutant and control lines, quantification of fluorescence images can be challenging as differences in the deposition of ECM components may be difficult to capture, even in extended cultures. Moreover, we only stained a limited number of components of the PNN. Therefore, we next aimed to expand our assessment of PNNs in our co-cultures by performing qPCR on a panel of genes including core PNN (*e.g.* versican *VCAN*, brevican *BCAN*, phosphacan *PTPRZ1*, *etc.*), and associated PNN regulatory genes (*e.g.* catabolic enzymes such as *MMP9*). We cultured MNs derived from iPSC with mutations in TDP-43 G298S, C9orf72 (C9-2), C9-3, and their controls (iso-TDP-43, iso-C9-2, and WT, respectively) and performed qPCR after 3, 6 and 9 weeks in culture using human-specific primers. We then plotted 2^dCt values for each gene and each cell line as heatmaps, and looked for differences in expression between mutants and their controls (Figures 2D-F and Table S3).

Using this approach, we found no significant differences in the expression of any genes when comparing C9-2 and C9-3 mutants to control lines at any time point. We did identify significant increases in expression of a handful of genes including *HAS3*, *HAPLN4*, *HYAL2* and *SEM3A* after 3 weeks culture when comparing MNs derived from TDP-43 G298S mutants to their isogenic control. Nevertheless, we also found increased expression of choline acetyl transferase (*CHAT*), a MN maturation marker^34^, in TDP-43 G298S MNs relative to isogenic controls, suggesting that increased PNN gene expression in TDP-43 G298S mutants might be attributable to faster maturation in these cells^36^. This interpretation was supported by a lack of differential regulation between these conditions at weeks 6 and 9. The lack of differential regulation between mutant and isogenic control lines could not be attributed to a loss of phenotype in culture as mutant lines showed significantly higher cytoplasmic TDP-43 localisation relative to their respective isogenic controls after 6 weeks in culture (Figure 2G-I and Supplementary Code 2). Therefore, we conclude that familial ALS mutations tested here do not affect the transcription of genes that form PNNs in iPSC-derived MNs cultured *in vitro*.

As we had only worked with a limited number of ALS mutant lines, we next aimed to apply our approach more broadly by screening the expression of genes encoding PNN components in additional ALS mutant lines by analysing 3^rd^-party data from a previously published meta-analysis^40^. Ziff *et al*. had analysed a range of transcriptomic datasets from ALS mutant iPSC-derived MNs, including sporadic cases and mutations in C9orf72, FUS, SOD1, and TARDBP. We generated a list of gene ontology (GO) terms for PNNs and calculated log fold change (LogFC) values and adjusted p-values comparing expression in mutant lines to that in MNs generated from control iPSC lines, which we then plotted as heat maps (Figure 2J). However, despite expanding the number of ALS-associated mutations that we analysed, we could not find strong evidence for consistent differential expression of genes associated with the formation of PNN components in mutants compared to controls. We did find that the gene for aggrecan (*ACAN*) was significantly upregulated in C9orf72 iPSC-derived MNs and that *PTPRZ1* was significantly downregulated in TARDBP mutant iPSC-derived MNs compared MNs derived from control iPSC lines. However, given that these were the only significantly differentially regulated genes across this large dataset, it seems unlikely that there is any intrinsic differential expression of PNN components by iPSC-derived MNs bearing ALS mutations.

### Altered expression of PNN components in human ALS post-mortem tissue is not captured by *in vitro* models

Although our *in vitro* model allowed us to create PNN-like structures from human cells, MNs containing ALS-associated mutations failed to show evidence of the loss of PNNs that have been reported in animal models at either the protein or transcriptomic levels. However, our *in vitro* cultures are reductionist and fail to incorporate many *in vivo* complexities, including the 3D environment, blood flow, and contributions from other neuronal and glial cell types. Therefore, we next assessed transcriptomic data captured from post-mortem human ALS tissue samples. Using bulk RNAseq data gathered in the same meta-analysis by Ziff et al.^40^, which included RNA from 214 ALS post-mortem spinal cord and 57 control samples, we then assessed the expression of PNN components in diseased tissues compared to controls. Using this approach, and contrary to our findings using iPSC-derived MNs, we found that multiple genes associated with PNN components were significantly differentially regulated in human post-mortem ALS tissues (Figure 3A). This included significant upregulation of *ACAN* and *PTPRZ1* (all cases), *VCAN* and *HAS2* (all except FUS) and *BCAN* (all except FUS and C9orf72), as well as significant down regulation of *HAPLN2* (all cases), *HAPLN3* (all except FUS) and *HAS3* (all except FUS and SOD1). Thus, these data point towards transcriptional regulation of PNN components in ALS tissue samples that are not captured in our *in vitro* model.

**Figure 3:**
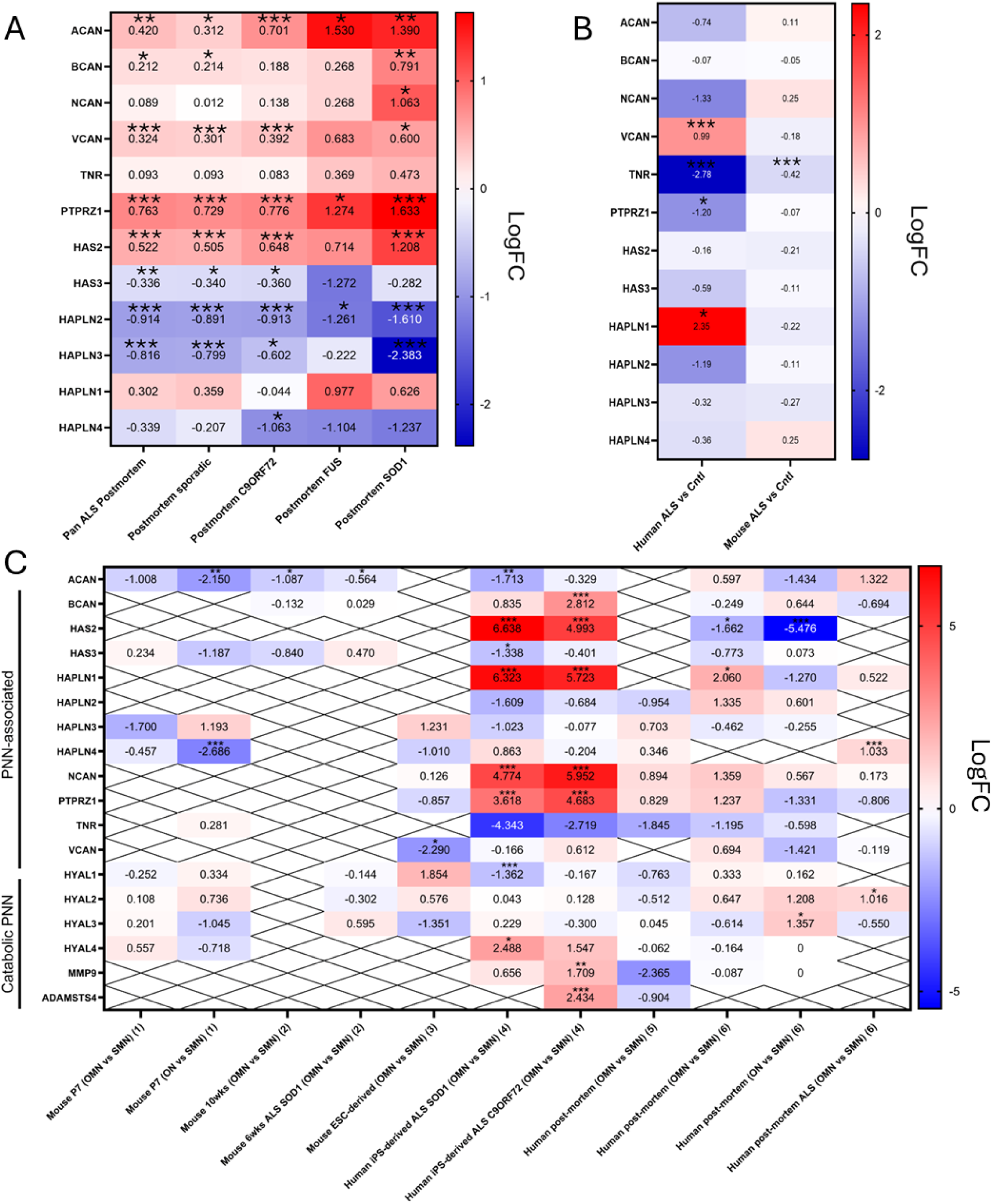
Altered expression of PNN components in human ALS post-mortem tissue is not captured by *in vitro* models. A) PNN expression in human post-mortem ALS tissues. A) Log fold change (LogFC)values of ALS mutant human post-mortem tissues for PNN component visualised as a heatmap. Relevant ALS mutation is indicated on x-axis. B) Log fold change (LogFC) values of ALS mutant human or mouse astrocytes for PNN related GO term visualised as a heatmap. Data acquired from Ziff et al. (2022). C) Expression of PNN and PNN-associated genes comparing resistant to susceptible tissues in healthy and ALS tissues and ALS cell models. Heatmaps show LogFC. Genes not expressed at detectable levels or that had multiple hits are indicted by an X through the relevant box. OMN = oculomotor neuron, ON= Onuf’s nucleus, SMN = spinal motor neurons. Each dataset is labelled by time point, model, comparison type (OMN vs SMN or ON vs SMN) and dataset origin (see Table S4). P-adjusted values, * <0.05, ** <0.01, *** <0.001, **** p < 0.0001

MNs are not the only cell type in the CNS that form PNNs. Indeed, glia such as astrocytes, microglia and oligodendrocytes also make PNN components^10^. Therefore, we again built from existing meta-analyses of astrocytes formed from hiPSC and mouse lines with ALS mutations^41^. In this analysis, mutations in human (SOD1, C9orf72, FUS) and mouse (SOD1 G93A mutation, *Tardbp* deletion, *Tmem259* deletion) lines were pooled together and corrected for batch effects. We then analysed differential expression of genes associated with PNN components by comparing to healthy controls (Figure 3B). We found that whilst mouse ALS astrocytes showed little differential regulation of genes for PNN components, for human iPSC lines with ALS mutations expression of *VCAN*, *TNR*, *PTPRZ1*, and *HAPLN1* were differentially regulated. Although upregulation of *VCAN* reflected changes we had observed in post-mortem tissues (Figure 3A), differential regulation of the other genes was not consistent. This suggested that although ALS mutations did have some impact on expression of PNN-associated genes in ALS mutant hiPSC astrocytes cultures, this alone was unlikely to account for the broad regulation of genes for PNN components we observed in human samples at end-stage disease.

In ALS, some subtypes of MNs are more susceptible to death than others, which can show selective resistance, even at the end stage of disease^6,7^. Indeed, ALS patients often maintain normal vision and can track an object with their eyes, but cannot speak, swallow, or move^6^. We failed to find evidence of a role for PNNs in ALS using iPSC-derived MNs, but observed significant regulation of gene expression for PNN components in human post-mortem tissues. This led us to hypothesise that our iPSC-derived MNs may not have been capturing the complexity of the selective resistance/vulnerability of MN populations *in vivo*. To address this, we next compiled a list of previously published studies that assessed differences in transcriptomic expression between MNs in susceptible and vulnerable areas to ALS using either bulk RNA sequencing or microarray (Table S4). We extracted expression levels from human and mouse data sets collected at various time points and disease states, and from different anatomical sites known to impart resistance or susceptibility in ALS (resistant: Onuf’s nucleus, ON or ocular motor neuron, OMN; vulnerable: spinal motor neuron, SMN). We then plotted logFC differences in genes encoding PNN components or their catabolic regulators comparing areas that are selectively vulnerable or resistant to MN death as a heat map (Figure 3C). Many PNN-associated genes were not expressed at detectable levels. Nevertheless, our analysis identified aggrecan (*ACAN*) as significantly downregulated between resistant (OMN) compared to vulnerable (SMN) tissues in certain datasets including 10-weeks healthy mouse, 6-weeks SOD1 mouse, and 9-day iPSC-derived SOD1 ALS mutant MNs. However, it was not significantly differentially expressed in any of the post-mortem human datasets, suggesting a lack of regulation at late-stages disease between vulnerable and resistant area.

However, when we assessed expression of PNN components in iPSC-derived OMNs and SMNs from SOD1 and C9orf72 ALS mutants, we found that OMNs derived from SOD1 mutant lines had significant differences in expression of *ACAN, HAS2, HAS3, HAPLN1, NCAN, PTPRZ1, HYAL1,* and *HYAL4* relative to SMNs. C9orf72 mutant line-derived OMNs similarly showed significant differences in expression of *BCAN, HAS2, HAPLN1, NCAN, PTPRZ1, MMP9,* and *ADAMSTS4* relative to SMNs. These data may indicate that PNN components are differentially regulated in OMN compared to SMN for C9orf72 and SOD1 ALS mutants. However, we did not observe similar changes in other datasets comparing resistant to vulnerable areas. Therefore, as these MNs were analysed only 9 days after the start of differentiation, and as we similarly observed some differential regulation of PNN components in TDP-43 G298S mutants that did not persist at later time points (Figure 2D-F), these differences might be attributable to differences in the development of the MNs themselves.

## Discussion

Primary neuron cultures from rodents have been shown to produce PNN-like structures *in vitro*^33,42,43^. However, to our knowledge, it has not been shown previously that human ESC-/iPSC-derived MNs can also create these ECM structures *in vitro*. To investigate if PNN-like structures were deposited by ESC-/iPSC-derived MNs, we established a co-culture system composed of wildtype human ESC-/iPSC-derived MNs and murine ESC-derived astrocytes. This system robustly generated PNN-like structures that stained positively for aggrecan, tenascin-R, and hyaluronan. iPSC-derived MNs become electrically mature between 2 and 6 weeks of culture^36^ – a similar time frame during which we observed deposition of PNN-like structures. These findings mirror those reported in *in vivo* models where initiation of electrical activity is reported to coincide with PNN formation^44^. Given that PNNs influence neuronal firing patterns^17^ and that hyperexcitability is a hallmark of ALS, our *in vitro* PNN-like model may offer a valuable platform to investigate how PNNs—and their loss—affect MN excitability in the context of ALS.

Following establishment of a culture system in which human ESC-/iPSC-derived MNs successfully form PNN-like structures, we next assessed the impact of ALS-associated mutations and found that the fraction of double positive MNs (β-III tubulin/PNN component: aggrecan, hyaluronan, tenascin-R) was no different in mutant lines compared to controls. Similarly, qPCR analysis of a panel of PNN-associated genes could not detect consistent differences in expression in mutant lines compared to controls, which we confirmed using data provided by a meta-analysis on iPSC-derived MNs formed from lines with additional ALS-associated mutations^40^. However, in post-mortem ALS tissues, the expression levels of multiple components of PNNs were altered. Indeed, we identified significant regulation of PNN components, including *ACAN*, *VCAN*, and *BCAN*, which were significantly upregulated, and *HAPLN2* and *HAPLN3*, which were significantly downregulated. Although it may seem counterintuitive that some core PNN components were upregulated while others were downregulated, this pattern may reflect complex compensatory responses. Alternatively, overlapping, uncoordinated responses across multiple cell types acting in parallel may yield mixed expression profiles. Moreover, post-mortem samples provide a snapshot of end-stage disease when many MNs have died, and the tissues are largely degenerated. In this context, the mechanisms that drove disease progression may no longer be at play. Indeed, it is possible that our iPSC-derived MNs might model an early phase of disease or not fully capture the complex disease process, which takes years to develop in humans.

Because PNNs are assembled and remodelled through contributions from multiple CNS cell types, the changes we observed in expression data taken from post-mortem tissues are likely shaped by cells that are not captured in our *in vitro* system. Glial cells in particular are central regulators of PNN formation and integrity. For example, in addition to MNs and astrocytes, oligodendrocytes can also supply key PNN components, including versican and link proteins^45^. Microglia that express MMP9 have also been identified surrounding denuded MNs and to even engulf the PNNs^21^. Moreover, α-MNs lacking PNNs show elevated expression of protein oxidation marker 3-nitrotyrosinen, a marker of protein oxidation and oxidative stress^21^. Given the established role of oxidative stress in ALS, these observations cautiously support a model in which microglial-mediated PNN degradation could compromise neuronal protection and increase oxidative stress, thereby contributing to α-MN’s vulnerability to cell death in ALS.

Kaplan and colleagues have shown previously that MMP9 expression is associated with areas of selective vulnerability to cell death in murine models of ALS, and that reducing MMP expression in these areas improved functional muscle output and increased survival times^28^. This highlights the value of comparing gene expression between resistant and vulnerable MN populations to identify candidates involved in selective susceptibility. To explore this, we performed a meta-analysis of publicly available datasets comparing resistant and vulnerable motor neurons in ALS. However, we could not find convincing evidence implicating PNNs in the resistance of ocular MNs. Although a few PNN-related genes were differentially expressed in resistant compared to susceptible tissues, many were not detected. Similarly, when we examined ALS astrocyte models, we observed only limited and inconsistent changes in PNN-related gene expression, with no clear patterns in either human or mouse models. This further supports the view that widespread PNN dysregulation in ALS is unlikely to be driven by cell-intrinsic differences in MNs or astrocytes alone. A more comprehensive transcriptomic analysis combined with functional studies will be needed to determine whether changes in PNN components contribute directly to neuroprotection or instead reflect regional differences in cell composition or activity.

Together, our findings demonstrate that human iPSC-derived MNs can form PNN-like structures *in vitro*, providing a cell culture model to study their function and regulation in ALS and other motor diseases. While our data suggest that MN-intrinsic mechanisms are not sufficient to explain PNN disruption in disease, changes in PNN-associated gene expression observed in post-mortem tissues point to potentially important roles for non-neuronal cells. This model lays the groundwork for future mechanistic studies into how cell-type interactions and matrix remodelling contribute to MN vulnerability and resistance in ALS.

## Resource availability

Materials availability: Cell lines are available on request and through appropriate MTAs.

Data and code availability: Data are either publicly available or are available upon request to the corresponding author. Code are available within the supplementary information included with this manuscript.

## Acknowledgments

CK acknowledges support from the Wellcome Trust Advanced Therapies for Regenerative Medicine PhD programme [218452/Z/19/Z] at King’s College London. EG and IL are grateful for support from the Rosetrees Trust. IL acknowledges support from the Medical Research Council [MR/N025865/1]. The authors wish the thank Dr Peter Harley and Dr Carolina Barcellos Machado for help with the cell lines. We also thank Dr Kevin Blighe for assistance with the bioinformatics work on the transcriptomic datasets comparing resistant and vulnerable tissues.

## Author contributions

C.K. performed all the experimental work and bioinformatics analyses. C.K., E.G. and I.L. designed the experiments. E.G. and I.L. supervised the project. C.K. and E.G. wrote the manuscript. All authors revised and approved the manuscript.

## Materials and Methods

### iPSC culture

Human induced pluripotent stem cells (iPSC) or human embryonic stem cells (ESC) (Table 1) were cultured in StemMacs™ iPSC brew supplemented with penicillin-streptomycin (1X) on laminin-521 basement matrix (LN521, Thermo Fisher, #A29249) diluted 1:50 (0.4 µg/cm^2^) in phosphate-buffered saline (PBS) containing calcium and magnesium (Thermo Fisher, #14040117). iPSC were passaged as single cells when ∼70% confluent. For passaging, confluent cells were detached from the culture plate by incubating with TrypLE express (Invitrogen) for 4 min at 37 °C. iPSC were then diluted in Dulbecco’s modified Eagle’s medium (DMEM) 1:10 (Thermo Fisher, #12491015) and centrifuged at 260 g for 4 min before plating the required number of cells in iPSC media supplemented with 10 µm Y-27632 (Tocris, Cat#1254) for 24 h. iPSC were maintained at 37°C with 5% CO_2_/95% air. Media was replenished every 24-48 h.

### Engineering of iPSC lines

To enable downstream purification of MNs and astrocytes, human iPSC and mESC were engineered to allow for cell type-specific magnetic activated cell sorting (MACS). The mESC line had been engineered previously with GFAP::CD14^35^. For MN differentiation, wildtype, TDP-43-G298S and Isogenic TDP-43-G298S human iPSC lines had all been previously engineered with Hb9::CD14^36^. To create C9orf72 mutant lines and their isogenic controls (C9-3, C9-2, iso-C9-2), we inserted the Hb9::CD14 (Addgene, #204344) construct via electroporation. 1 µg/µl plasmid DNA was added in OptiMEM Reduced-Serum Medium (Gibco, 11058021). iPSC were detached from LN521 coated plates using TrypLE express, washed once with DMEM, and then twice with Opti-MEM Reduced Serum Medium™ before concentrating cells to 1 million iPSC/90µl. For Hb9::CD14 insertion into the AAVS1 (Adeno-associated virus integration site 1) locus, 5 µg Hb9::CD14 plasmid, 2.5 µg TALEN-Left (Addgene #52341) and 2.5 µg TALEN-Right (Addgene #52342) were mixed with iPSC. Each iPSC/plasmid mixture was added to 2 mm electroporation cuvettes (GeneFlow, E6-0062) and electroporated. Post-electroporation, iPSC were collected and replated in LN521-coated 60 mm dishes in pre-warmed iPSC media supplemented with 10 µm Y-27632 for the first 24 h, and were allowed to reach 70% confluency. iPSC were then treated with 0.5 µg/µl puromycin for 5 days for selection of transfected cells. After recovery, single iPSC were plated by limiting dilution into LN521-coated 96-well plates (0.5 cells/well). Surviving iPSC colonies were dissociated using TrypLE and genomic DNA was isolated using QuickExtract (Cambio/Epicentre, #QE09050^48^). Genomic DNA was amplified by PCR and screened for homozygous insertion of the Hb9::CD14 plasmid.

### Generation of motor neurons (MNs) from iPSC lines

MNs were generated from iPSC lines as described previously^34^. Single iPSC were plated in suspension culture plates in MN differentiation media (N.MND, Table 2) supplemented with 10 μM Y-27632, 20 μM SB431542, 0.1 μM LDB193189, and 3 μM CHIR. On day 2, media was replaced with N.MND media supplemented as above with the addition of 100 nM RA and 500 nM SAG. On day 4 media was replaced with N.MND media supplemented with 20 μM SB431542, 0.1 μM LDB193189, 100 nM RA and 500 nM SAG. On day 7 media was changed to N.MND media supplemented with 100 nM RA and 500 nM SAG. On day 9 media was changed to N.MND media supplemented with 500 μM ascorbic acid, 0.1 μM compound E, 100 nM RA and 500 nM SAG. On day 11 media was changed to N.MND media supplemented with 500 μM ascorbic acid, 0.1 μM compound E, 100 nM RA, 10 ng/mL BDNF, 10 ng/mL GDNF, 10 ng/mL CNTF, 10 ng/mL NT3, 10 ng/mL IGF1, 10 μM forskolin, 100 μM IBMX and 500 nM SAG. On day 14, MNs were purified using MACS.

**Table 2:** Composition of N.MND.

| Component | Volume in 50mL | Manufacturer |
| --- | --- | --- |
| Advanced DMEM/F12 | 24mL | Thermo Fisher Scientific |
| Neurobasal media | 24mL | Thermo Fisher Scientific |
| Penicillin-streptomycin, 100X | 0.5mL | Thermo Fisher Scientific |
| L-Glutamine (200mM) | 0.5mL | Thermo Fisher Scientific |
| NeuroBrew-21 supplement without vitamin A (50X) | 0.5mL | Miltenyi Biotech |
| N-2supplement | 0.25mL | Thermo Fisher Scientific |
| β-mercaptoethanol1000x | 200µL | Thermo Fisher Scientific |

### Generation of astrocytes from mESC

The mouse ESC E14-IB10 subclone G6 was previously engineered to express GFAP::CD14/CAG::GDNF^37^. mESC were cultured in mESC maintenance media (Table 3) on LN521-coated plates (0.08 μg/cm^2^) until 70% confluent, with maintenance media replaced every 1-2 days. 1 million mESC were placed in 10 cm^2^ suspension plates with astrocyte media (Table 4) for 2 days. On day 2, EBs were collected, split 1:4 and then cultured in astrocyte media supplemented with 0.5 μM SAG and 1 μM RA for 3 days. On day 5, EBs were collected and replated in astrocyte media in T175 flasks coated with Matrigel (1:100 diluted in DMEM) and cultured in astrocyte media for a further 7 days until purification by MACS.

**Table 3:** Composition mESC maintenance medium.

| Component | Volume in 50mL | Manufacturer |
| --- | --- | --- |
| DMEM/F12 Medium (1X) | 22.5mL | Thermo Fisher Scientific |
| Neurobasal media | 22.5mL | Thermo Fisher Scientific |
| Penicillin-streptomycin, 100X | 0.5mL | Thermo Fisher Scientific |
| L-Glutamine (200mM) | 0.5mL | Thermo Fisher Scientific |
| MACS NeuroBrew-21 without vit A (50X) | 0.5mL | Miltenyi Biotech |
| N-2 supplement | 0.25mL | Thermo Fisher Scientific |
| β-mercaptoethanol1000x | 50µL | Thermo Fisher Scientific |
| PD0325901 | 5µL | Tocris |
| CHIR99021 | 15µL | Tocris |
| Plasmocin Prophylactic (25mg/ml) | 10µL | InvivoGen |
| 2 inhibitor/leukemia inhibitory factor (2iLIF) supernatant (COS7/pCAGGS::LIF) | 0.5mL |  |
| ESC-qualified Fetal BSA | 1mL | Gibco |
| 5% Sterile BSA fraction V | 1mL | Roche |

**Table 4:** Composition of astrocyte medium.

| Component | Volume in 50mL | Manufacturer |
| --- | --- | --- |
| Advanced DMEM/F12 | 24mL | Thermo Fisher Scientific |
| Neurobasal media | 24mL | Thermo Fisher Scientific |
| Penicillin-streptomycin, 100X | 0.5mL | Thermo Fisher Scientific |
| L-Glutamine (200mM) | 0.5mL | Thermo Fisher Scientific |
| NeuroBrew-21 supplement without vitamin A (50X) | 0.5mL | Miltenyi Biotech |
| N-2supplement | 0.25mL | Thermo Fisher Scientific |
| β-mercaptoethanol1000x | 200µL | Thermo Fisher Scientific |
| 5% Sterile BSA fraction V | 1mL | Roche |

### Magnetic activated cell sorting (MACS) purification of MNs and astrocytes

MACS purification of MNs was performed as described previously^36^. Briefly, day 14 EBs were dissociated using trypsin-EDTA supplemented with 10 U/mL DNase-I (Merck, #4536282001) for 8-10 mins. DMEM supplemented with 10 U/mL DNase-I and 10% FBS (Thermo Fisher, #26140079) was then added to the EB/trypsin-EDTA mixture and centrifuged at 500 rpm for 4 mins and supernatant aspirated. To obtain single cells, EBs were resuspended in 500 µL DMEM + DNase-I (10 U/mL) and were mechanically disrupted by pipetting up and down 10-12 times. 1 mL DMEM + DNase-I (10U/mL) was then added and remaining EBs allowed to settle for 5 mins. 1 mL of supernatant was removed and placed in a new collection tube, and this process was repeated 3 times. EB supernatant solution was then passed through a 40 µm strainer, centrifuged and washed once in MACS buffer (90% PBS, 10% 0.5% BSA and 10 U/mL DNase-I). 3 µg/mL anti-CD14 diluted in MACS buffer was added to the cell suspension and incubated on a rotator for 15 min at 4 °C. Cells were washed once with MACS buffer before adding anti-mouse IgG microbeads (1:10, Miltenyi, #130-048-402) and the solution was again incubated on a rotator for 20 min at 4 °C. Following an additional wash in MACS buffer, cells were resuspended in MACS buffer before passing through a MACS column (Miltenyi, #130-042-201) that had been pre-washed with MACS buffer. Cells retained on the column were then resuspended in N.MND media. To purify mESC-derived astrocytes, a similar procedure was used with the exception that as astrocyte cultures were adherent and grown in T175 flasks, trypsin-EDTA supplemented with 10 U/mL DNase-I (15 mins) was used to detach cells before being passed through a 40 µm strainer. Eluted cells were resuspended in astrocyte media.

### Establishment of iPSC-derived MNs and mESC-derived astrocyte co-cultures

Purified populations of MNs and astrocytes were used to generate co-cultures. Astrocytes were plated in astrocyte media 5-7 days before MNs were added to allow recovery and cell spreading, before plating MNs on top. For co-cultures, a 50:50 mixture of astrocyte medium:N.MND media was used with 10 µm Y-27632 added for the first 24 h of culture.

### qPCR analysis of MNs-astrocyte co-cultures

Total RNA was isolated from cells using an RNeasy Mini Kit (Qiagen, 74104), following the manufacturer’s guidelines, and RNA quantified using a NanoDrop 2000 (Thermo Fisher Scientific). Single stranded cDNA was generated using a High-Capacity cDNA Reverse Transcription Kit (Thermo Fisher Scientific, #4368814) using the manufacturer’s guidelines. 6.25 ng/µL cDNA in a final volume of 10 µL was used for qPCR. A mixture of 50% Fast SYBR Green PCR Master Mix (Thermo Fischer Scientific, 4385612), 35% water (Thermo Fisher Scientific, #AM9935) and oligonucleotide primers (2.5% forward and 2.5% reverse primer) (Sigma-Aldrich, Table 5) was prepared for each primer pair. This mixture was added to each well with 1µL cDNA solution and qPCR performed on a CFX384 Touch Real-Time PCR Detection System (BioRad) at the following settings: 95 °C 10 min, 40 cycles of 95 °C for 30 s and 60 °C for 1 min, 75 °C for 10 s, 95 °C for 10 s. Housekeeping genes and data was analysed either by -ΔCt (threshold cycle value) or Comparative CT Method (2ΔCt) using *ACTB* and *GAPDH* as housekeeping genes. Efficiency tests were performed on primers using human spinal cord total RNA (Takara bio, #636554), which confirmed that all had efficiencies between 85 and 105%.

**Table 5:**
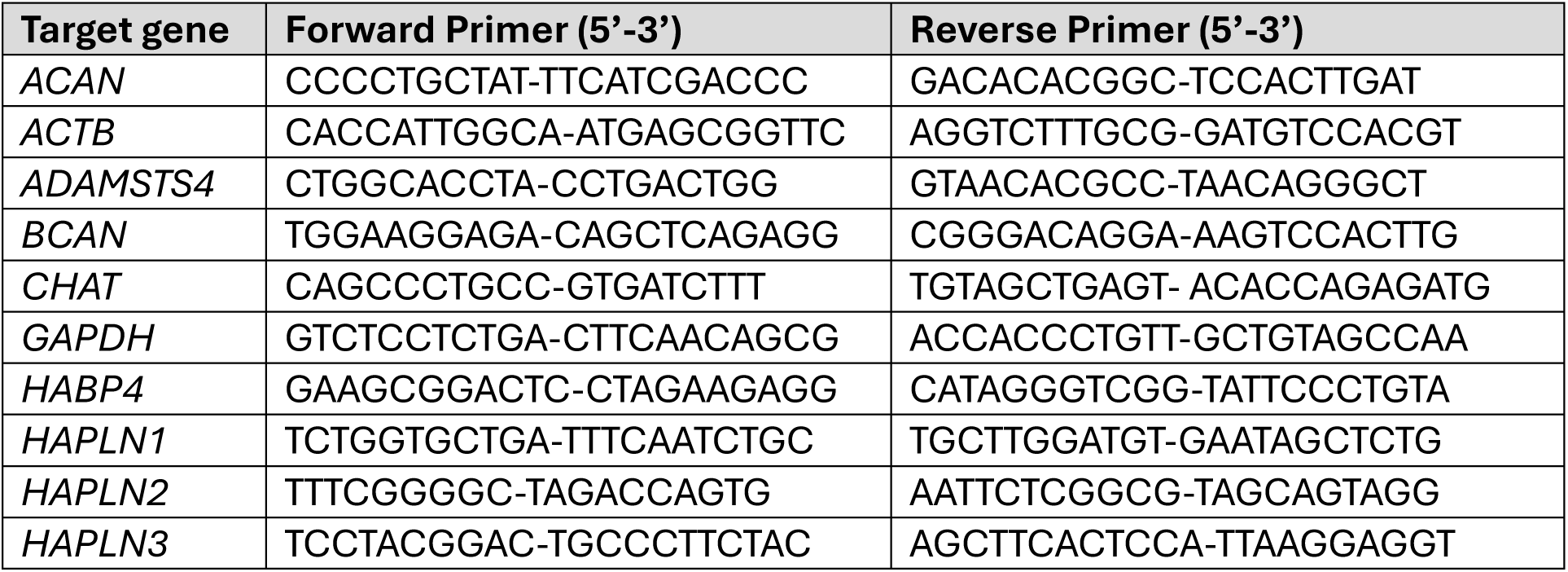
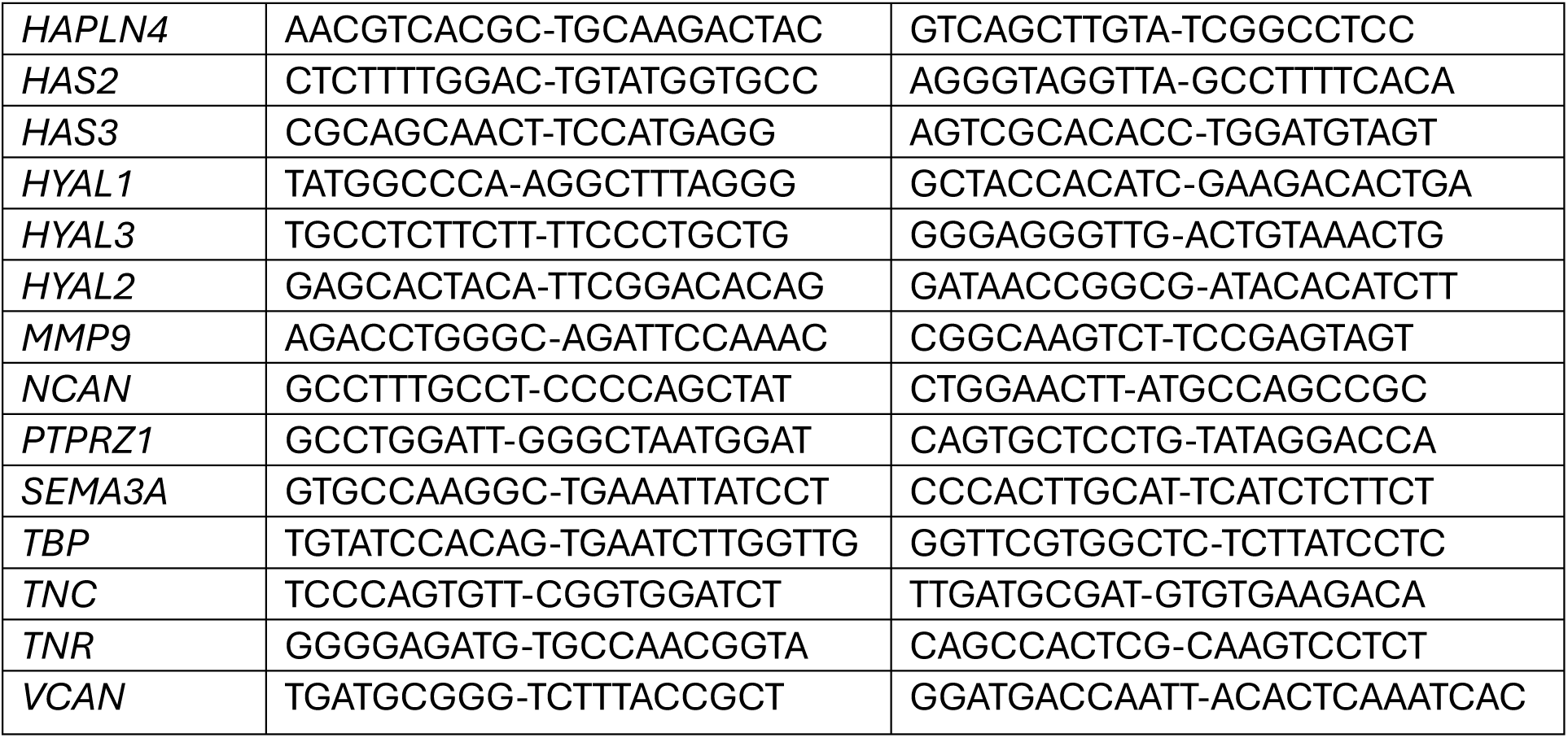
Primers used for qPCR.

### Immunocytochemistry staining of co-cultures

Half the volume of media was removed from wells and replaced with 4% w/v paraformaldehyde (PFA) (Thermo Fisher Scientific, 11586711) supplemented with 15% sucrose. After 10 min, half of this solution was removed, and another fresh volume of PFA/sucrose was added to the well for another 10 min. The total volume was then removed and the well washed twice with PBS. A combination permeabilisation-blocking step was then performed with 3% v/v Horse Serum (Sigma-Aldrich, #H0146) in PBS-T (0.1% Triton X-100 (Thermo Fisher Scientific) in PBS) for 30 min. Cultures were incubated overnight at 4 °C with primary antibodies diluted as indicated in Table 6. The following day, wells were washed three times with DPBS before adding secondary antibodies. Secondary antibody incubation occurred at room temperature for 45 mins before washing 3 times with DPBS. Samples were then imaged using either a Leica TCS SP8 (Leica Microsystems) or Zeiss LSM 980.

**Table 6:** Antibodies used for immunostaining.

| **Antibody** | **Catalogue number** | **Species** | **Supplier** | **Concentration** |
| --- | --- | --- | --- | --- |
| Aggrecan core protein | bs-1223R-BSS | Rabbit | Bioss | 1:200 |
| Aggrecan | Ab1031 | Rabbit | Merck | 1:200 |
| Aggrecan | A279519 | Mouse | Antibodies online | 1:200 |
| Aggrecan | MA316888 | Mouse | Thermo Fisher | 1:200 |
| Hyaluronic acid binding protein | 385911 | Biotinylated | Sigma Aldrich | 1:200 |
| hTenascin R pAb | AF3865-SP | Goat | RnD Systems | 1:200 |
| Wisteria floribunda agglutinin | L1516 | Biotinylated | Sigma Aldrich | 1:200 |
| Beta-III tubulin | MAB1195 | Mouse | RnD Systems | 1:1000 |
| Choline Acetyl-transferase | AB114P | Goat | Sigma Aldrich | 1:200 |
| Choline Acetyl-transferase | NBP2-46620 | Mouse | Novus Biologics | 1:200 |
| TDP-43 | 107-82-2-AP | Rabbit | Protein Tech | 1:300 |
| Streptavidin AlexaFluor™ 647 | S32357 |  | Abcam | 1:1000 |
| Donkey anti-Mouse IgG, AlexaFluor™ 488 | A-21202 | Donkey | Thermo Fisher | 1:1000 |
| Donkey anti-Mouse IgG, AlexaFluor™ 546 | A10036 | Donkey | Thermo Fisher | 1:1000 |
| Donkey anti-Mouse IgG2a, AlexaFluor™ 647 | A-21241 | Goat | Thermo Fisher | 1:1000 |
| Donkey anti-Rabbit IgG, AlexaFluor™ 488 | A21126 | Donkey | Thermo Fisher | 1:1000 |
| Donkey anti- Rabbit IgG, AlexaFluor™ 568 | A10042 | Donkey | Thermo Fisher | 1:1000 |
| Donkey anti-Rabbit IgG, AlexaFluor™ 647 | A21050 | Donkey | Thermo Fisher | 1:1000 |
| Donkey anti-Goat IgG, AlexaFluor™ 568 | A-11057 | Donkey | Thermo Fisher | 1:1000 |
| Donkey anti-Rat IgG, AlexaFluor™ 568 | A78946 | Donkey | Thermo Fisher | 1:1000 |
| Donkey anti-Rat IgG, AlexaFluor™ 647 | A48272 | Donkey | Thermo Fisher | 1:1000 |

### Statistical analysis

Unless otherwise stated, data are presented as mean +/− standard error of the mean (SEM). Details of each test used are given in the relevant figure legends. Data were analysed using Graphpad Prism, Version 10 (Graphpad Software, San Diego, California USA).

---

**Figure S1:**
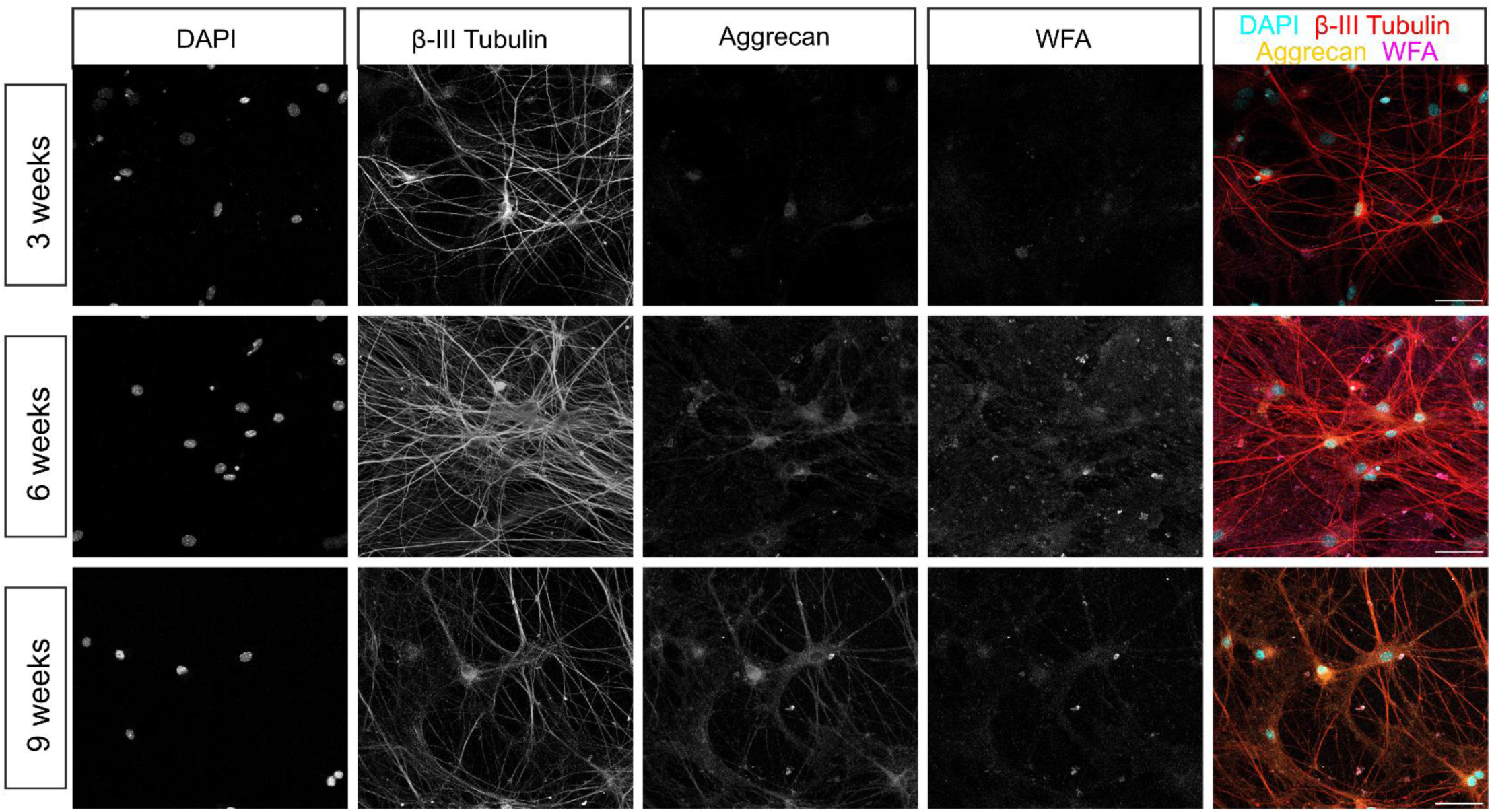
Representative confocal microscopy images of 3-, 6- and 9-week co-cultures of human ESC-derived MNs and astrocytes. Staining for β-III tubulin, aggrecan, Wisteria Floribunda Agglutin (WFA) and DAPI. WFA staining highlights similar structures to antibodies against aggrecan. Scale bar = 25μm.

**Figure S2:**
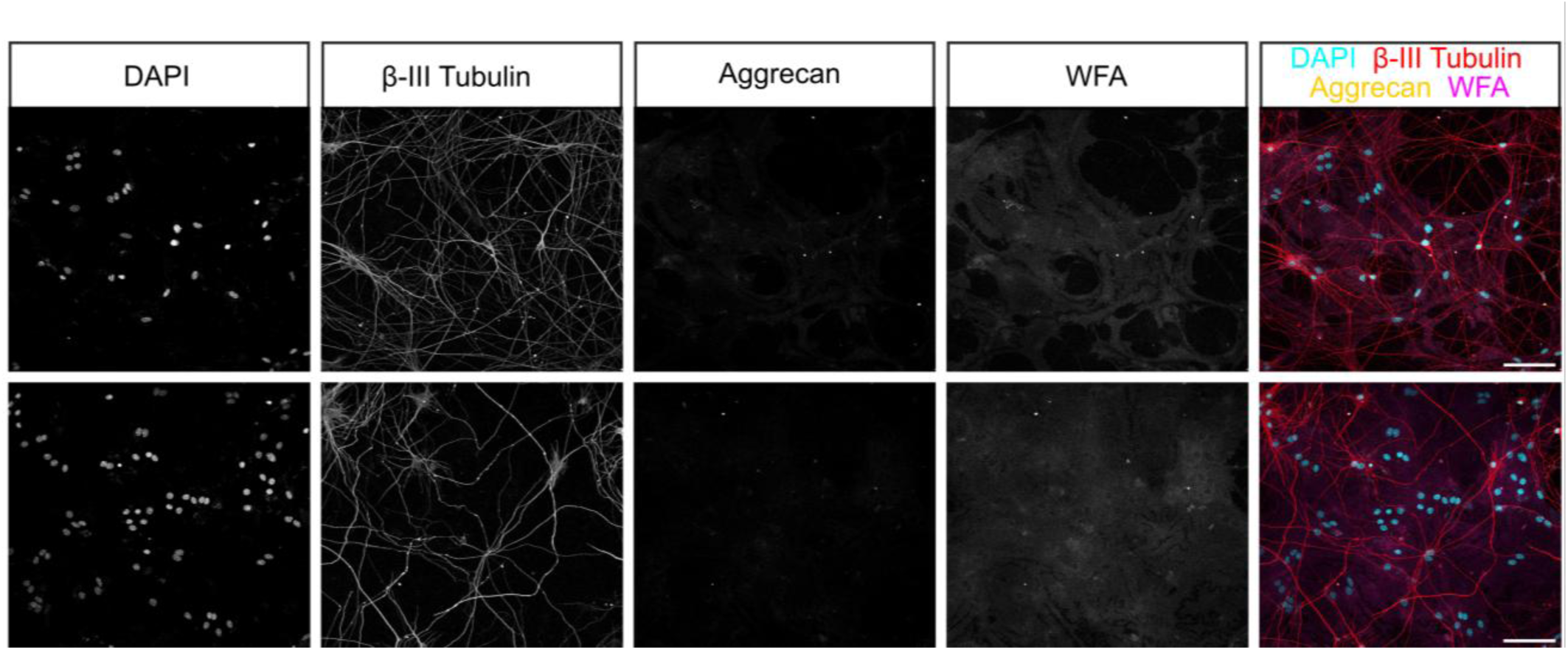
Representative confocal microscopy images of 2-week co-cultures of WT human ESC-derived MNs and astrocytes stained for β-III tubulin, aggrecan, Wisteria Floribunda Agglutin (WFA) and DAPI. PNN-like structures are not apparent at this stage. Scale bar = 100µm.

**Figure S3:**
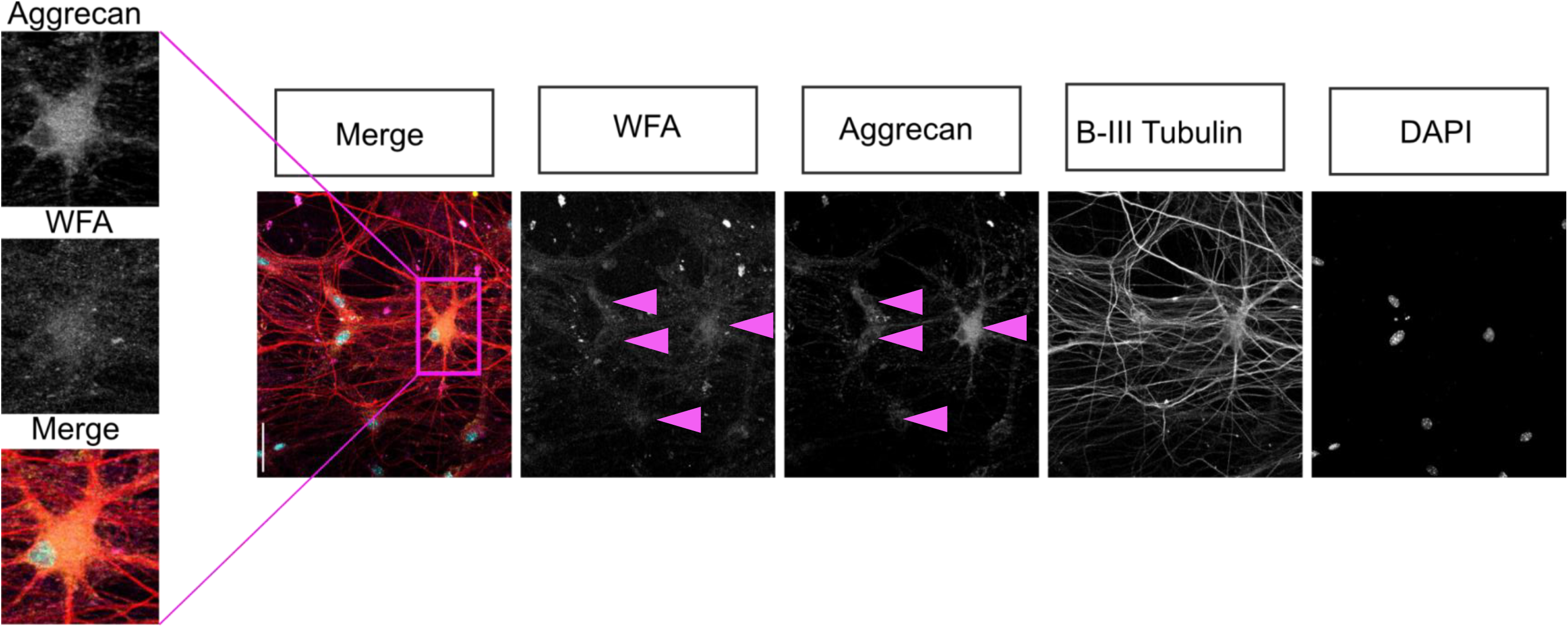
Representative confocal microscopy images of 9-week co-cultures of WT human ESC-derived MNs and astrocytes stained for β-III tubulin, aggrecan, Wisteria Floribunda Agglutin (WFA) and DAPI. Pink arrowheads point towards areas corresponding to β-III tubulin positive MNs. Scale bar = 50μm. Images acquired on Leica SP8 scanning laser confocal microscope.

**Figure S4:**
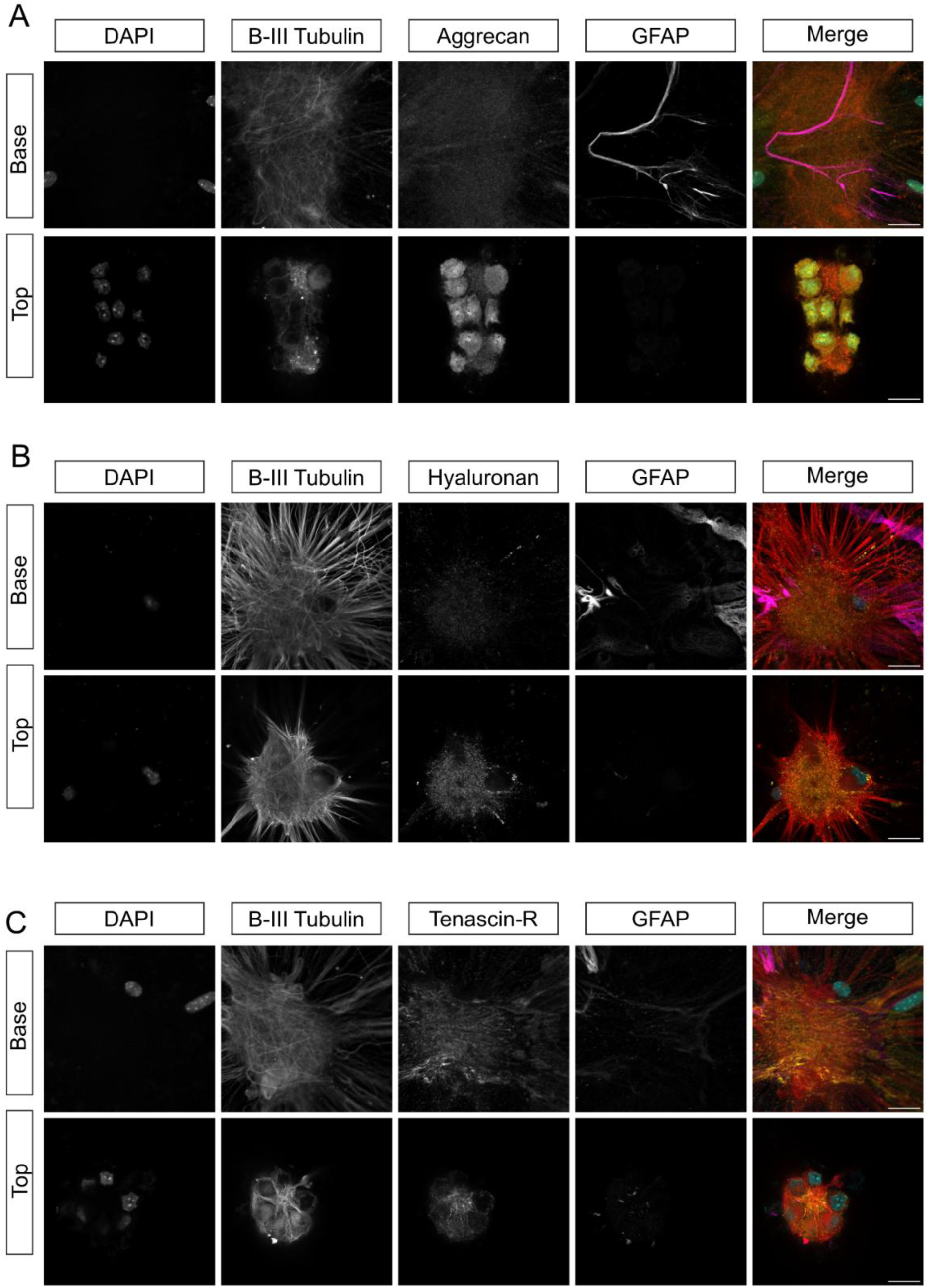
PNN components do not accumulate around astrocytes. Representative confocal microscopy images captured after 9 weeks of culture showing staining for PNN components A) aggrecan, B) hyaluronan, and C) tenascin-R captured from both the base and top of the cultures. Staining for PNN components almost never co-localised with GFAP+ astrocytes (GFAP+) which were present at the base of co-cultures. PNN instead accumulated around β-tubulin+ MNs. Scale bar = 25µm.

**Figure S5:**
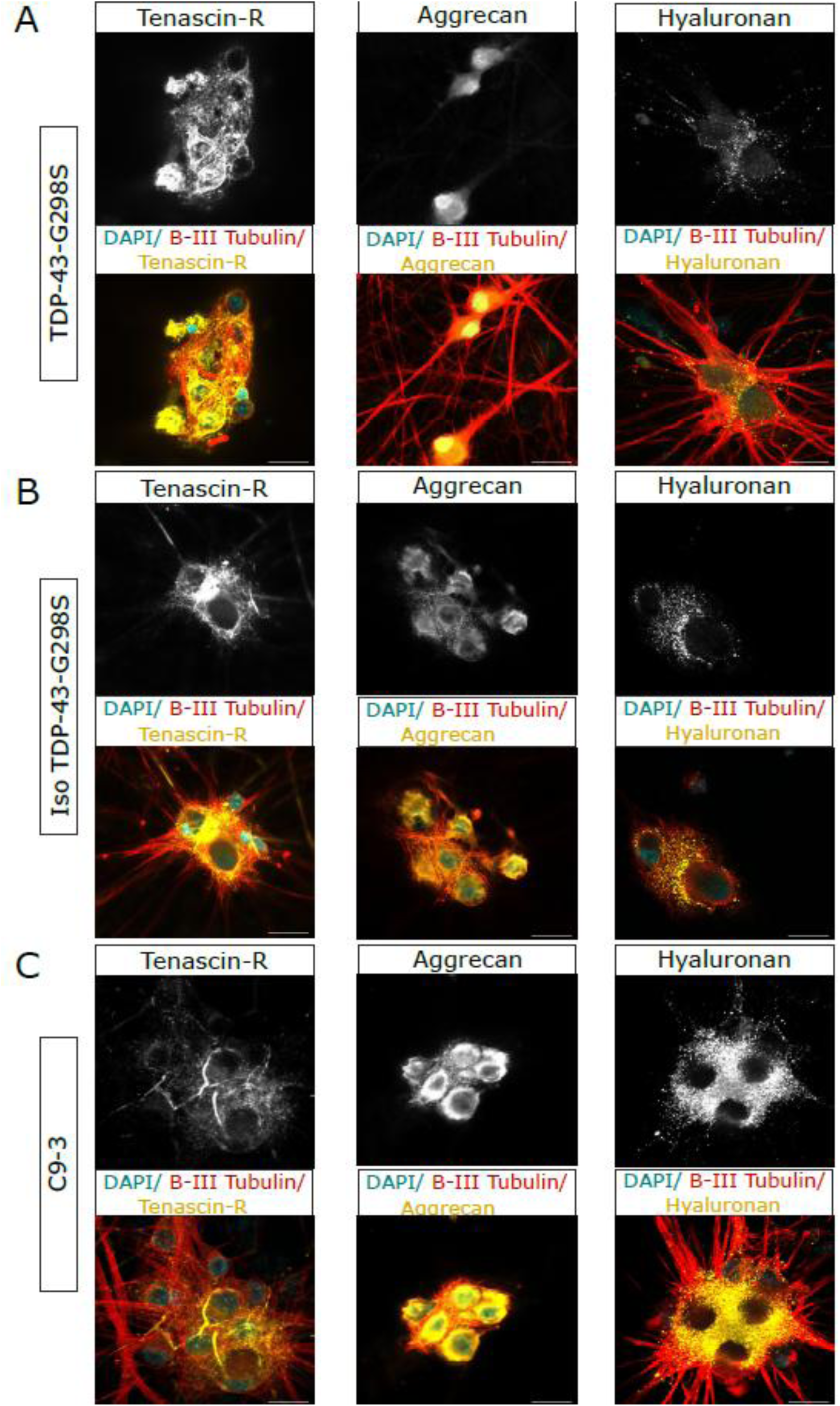
Representative confocal microscopy images of co-cultures stained for β-III tubulin, aggrecan, hyaluronan and tenascin-R after 9 weeks in culture. Scale bar = 25μm. Zeiss LSM confocal microscope, 63X. Representative images from 2 healthy lines (H9 and Iso-TDB-43-G298S) and two ALS line (C9-3 and TDB-43-G298S) all produce PNN components robustly.

**Table S1:** Summary of statistical tests used to assess significant differences between double positive β-III tubulin/PNN component (aggrecan/hyaluronan/tenascin-R) MNs in 3, 6 or 9 week co-cultures of WT ESC-derived MNs and astrocytes

| Hyaluronan |  |  |  |  |
| --- | --- | --- | --- | --- |
| Tukey's multiple comparisons test | Mean Diff. | Below threshold? | Summary | Adjusted P Value |
| 3wks vs. 6wks | -32.35 | Yes | ** | 0.0047 |
| 3wks vs. 9wks | -42.02 | Yes | ** | 0.0012 |
| 6wks vs. 9wks | -9.667 | No | ns | 0.3304 |
| Aggrecan |  |  |  |  |
| Tukey's multiple comparisons test | Mean Diff. | Below threshold? | Summary | Adjusted P Value |
| 3wks vs. 6wks | -44.74 | Yes | * | 0.0247 |
| 3wks vs. 9wks | -26.95 | No | ns | 0.1488 |
| 6wks vs. 9wks | 17.78 | No | ns | 0.3747 |
| Tenascin-R |  |  |  |  |
| Tukey's multiple comparisons test | Mean Diff. | Below threshold? | Summary | Adjusted P Value |
| 3wks vs. 6wks | -54.97 | Yes | * | 0.0204 |
| 3wks vs. 9wks | -71.94 | Yes | ** | 0.0059 |
| 6wks vs. 9wks | -16.97 | No | ns | 0.5056 |

**Table S2:** Statistical details for analysis of PNN-like deposition ICC for each iPSC-derived MN line and wildtype ESC derive MN line after co-culture of 3, 6 or 9 weeks. . Analysis performed was a one-way ANOVA with Tukey’s multiple comparison tests. 5 images/N (N=3) were counted for percentage of double positive MNs (β-III tubulin and hyaluronan/tenascin-R/aggrecan).

| Tenascin-R |  |  |  |  |  |
| --- | --- | --- | --- | --- | --- |
|  | Tukey's multiple comparisons test | Mean Diff. | Below threshold? | Summary | Adjusted P Value |
| 3 weeks | WT vs. C9-3 | -7.556 | No | ns | 0.9864 |
|  | WT vs. TDP-43 | 5.25 | No | ns | 0.9953 |
|  | WT vs. Iso-TDP-43 | -14.75 | No | ns | 0.9114 |
|  | C9-3 vs. TDP-43 | 12.81 | No | ns | 0.9394 |
|  | C9-3 vs. Iso-TDP-43 | -7.194 | No | ns | 0.9882 |
|  | TDP-43 vs. Iso-TDP-43 | -20 | No | ns | 0.8083 |
| 6 weeks | WT vs. C9-3 | 16.37 | No | ns | 0.8837 |
|  | WT vs. TDP-43 | 6.111 | No | ns | 0.9927 |
|  | WT vs. Iso-TDP-43 | 1 | No | ns | >0.9999 |
|  | C9-3 vs. TDP-43 | -10.25 | No | ns | 0.9674 |
|  | C9-3 vs. Iso-TDP-43 | -15.37 | No | ns | 0.9013 |
|  | TDP-43 vs. Iso-TDP-43 | -5.111 | No | ns | 0.9957 |
| 9 weeks | WT vs. C9-3 | 24.49 | No | ns | 0.6964 |
|  | WT vs. TDP-43 | 32.02 | No | ns | 0.4935 |
|  | WT vs. Iso-TDP-43 | 16.97 | No | ns | 0.8723 |
|  | C9-3 vs. TDP-43 | 7.526 | No | ns | 0.9866 |
|  | C9-3 vs. Iso-TDP-43 | -7.519 | No | ns | 0.9866 |
|  | TDP-43 vs. Iso-TDP-43 | -15.04 | No | ns | 0.9066 |
| Hyaluronan |  |  |  |  |  |
|  | Tukey's multiple comparisons test | Mean Diff. | Below threshold? | Summary | Adjusted P Value |
| 3 weeks | WT vs. C9-3 | 24.43 | No | ns | 0.1644 |
|  | WT vs. TDP-43 | 2.399 | No | ns | 0.9965 |
|  | WT vs. Iso-TDP-43 | 4.315 | No | ns | 0.9807 |
|  | C9-3 vs. TDP-43 | -22.03 | No | ns | 0.2368 |
|  | C9-3 vs. Iso-TDP-43 | -20.11 | No | ns | 0.3092 |
|  | TDP-43 vs. Iso-TDP-43 | 1.915 | No | ns | 0.9982 |
| 6 weeks | WT vs. C9-3 | 12.4 | No | ns | 0.6962 |
|  | WT vs. TDP-43 | 0.05555 | No | ns | >0.9999 |
|  | WT vs. Iso-TDP-43 | -4.944 | No | ns | 0.9716 |
|  | C9-3 vs. TDP-43 | -12.34 | No | ns | 0.6991 |
|  | C9-3 vs. Iso-TDP-43 | -17.34 | No | ns | 0.4355 |
|  | TDP-43 vs. Iso-TDP-43 | -5 | No | ns | 0.9706 |
| 9 weeks | WT vs. C9-3 | 5 | No | ns | 0.9706 |
|  | WT vs. TDP-43 | 1.905 | No | ns | 0.9983 |
|  | WT vs. Iso-TDP-43 | -2 | No | ns | 0.998 |
|  | C9-3 vs. TDP-43 | -3.095 | No | ns | 0.9927 |
|  | C9-3 vs. Iso-TDP-43 | -7 | No | ns | 0.9253 |
|  | TDP-43 vs. Iso-TDP-43 | -3.905 | No | ns | 0.9856 |
| Aggrecan |  |  |  |  |  |
|  | Tukey's multiple comparisons test | Mean Diff. | Below threshold? | Summary | Adjusted P Value |
| 3 weeks | WT vs. C9-3 | -23.45 | No | ns | 0.7057 |
|  | WT vs. TDP-43 | -10.89 | No | ns | 0.9581 |
|  | WT vs. Iso-TDP-43 | -10.29 | No | ns | 0.9643 |
|  | C9-3 vs. TDP-43 | 12.56 | No | ns | 0.9378 |
|  | C9-3 vs. Iso-TDP-43 | 13.17 | No | ns | 0.9294 |
|  | TDP-43 vs. Iso-TDP-43 | 0.6032 | No | ns | >0.9999 |
| 6 weeks | WT vs. C9-3 | -0.06611 | No | ns | >0.9999 |
|  | WT vs. TDP-43 | 7.414 | No | ns | 0.986 |
|  | WT vs. Iso-TDP-43 | 8.486 | No | ns | 0.9794 |
|  | C9-3 vs. TDP-43 | 7.48 | No | ns | 0.9857 |
|  | C9-3 vs. Iso-TDP-43 | 8.552 | No | ns | 0.9789 |
|  | TDP-43 vs. Iso-TDP-43 | 1.072 | No | ns | >0.9999 |
| 9 weeks | WT vs. C9-3 | 19.75 | No | ns | 0.8008 |
|  | WT vs. TDP-43 | 5.635 | No | ns | 0.9937 |
|  | WT vs. Iso-TDP-43 | -10.67 | No | ns | 0.9605 |
|  | C9-3 vs. TDP-43 | -14.11 | No | ns | 0.9149 |
|  | C9-3 vs. Iso-TDP-43 | -30.41 | No | ns | 0.5125 |
|  | TDP-43 vs. Iso-TDP-43 | -16.3 | No | ns | 0.876 |

**Table S3:**
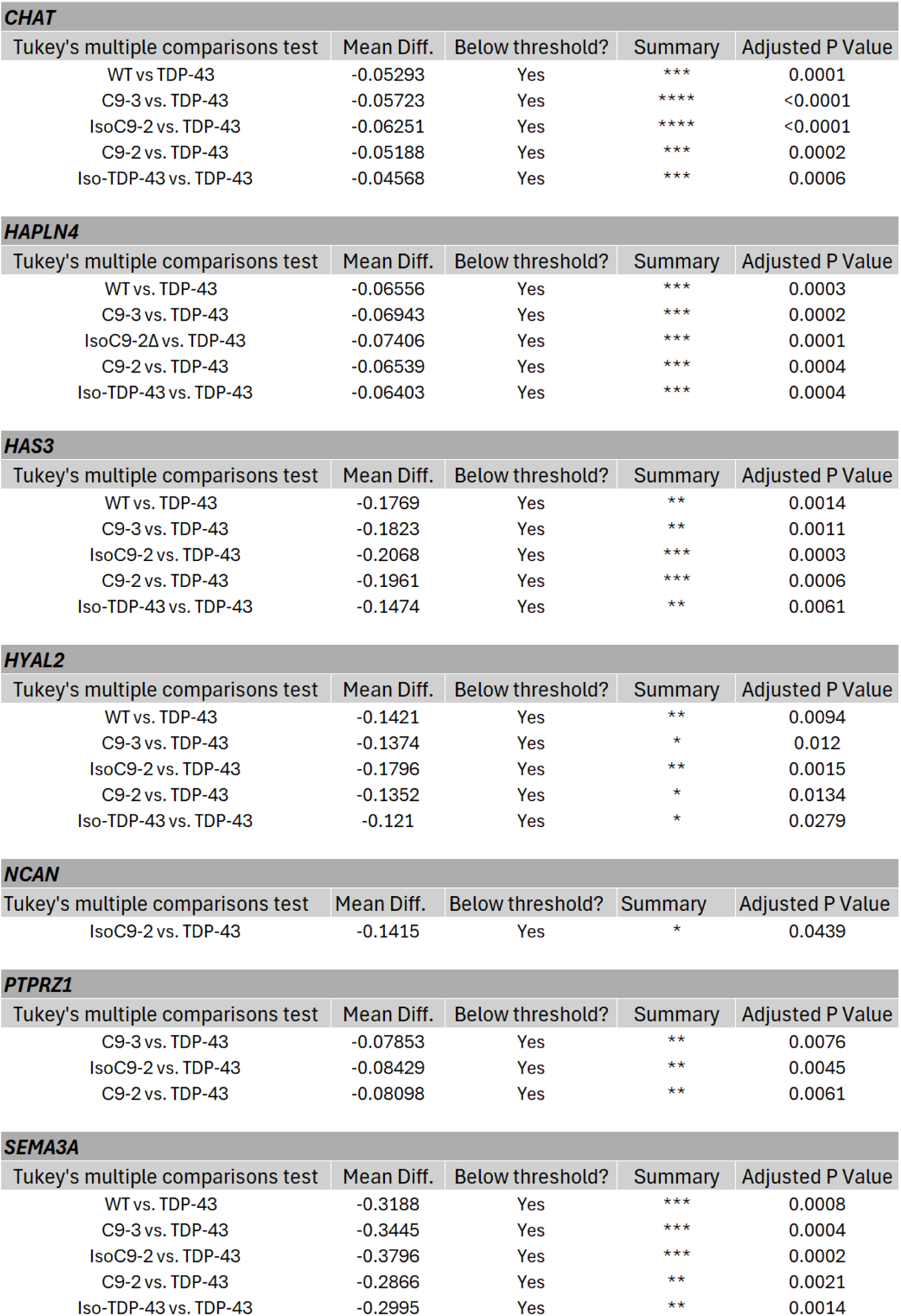
Details of statistical tests performed on qPCR data generated for hESC/hiPSC derived MNs after 3 weeks of co-culture with mouse astrocytes. Only details of significantly differentially expressed genes are given here.

**Table S4:** Datasets used in transcriptomic analysis to compare PNN gene expression in vulnerable and resistant tissues. Abbreviations: OMN: oculomotor nerve/cranial nerve III, SMN: spinal cord MNs, mESC: mouse embryonic stem cells, P7: postnatal day 7, hiPSC: human induced pluripotent stem cells.

| Dataset number | GEO ascension number | Details | Species | Age | Disease status | Anatomical area | Direction of comparison |
| --- | --- | --- | --- | --- | --- | --- | --- |
| 1 | E-GEOD_52118 | Microarray, laser capture microdissection | Mouse | P7 | Healthy/non-ALS | OMN | OMN vs SMN |
| 1 | E-GEOD_52118 | Microarray, laser capture microdissection | Mouse | P7 | Healthy/non-ALS | Onuf's nucleus | Onuf's nucleus vs SMN |
| 2 | GSE3343 | Microarray | Mouse | 10 wks | Healthy/non-ALS | OMN | OMN vs SMN |
| 2 | GSE3343 | Microarray | Mouse | 6 wks | ALS - SOD1 | OMN | OMN vs SMN |
| 3 | GSE118620 | Bulk RNA sequencing | mESC | 9 days in culture | Healthy/non-ALS | OMN | OMN vs SMN |
| 4 | GSE132972 | Bulk RNA sequencing | hiPSC | 17 days in culture | ALS - SOD1 | OMN | OMN vs SMN |
| 4 | GSE132972 | Bulk RNA sequencing | hiPSC | 17 days in culture | ALS - C9ORF72 | OMN | OMN vs SMN |
| 5 | E-GEOD-40438 | Microarray, laser capture microdissection | Human | Post-mortem, age 59 | Healthy/non-ALS | OMN | OMN vs SMN |
| 6 | GSE93939 | Bulk RNA sequencing; laser capture microdissection | Human | Post-mortem | Healthy/non-ALS | OMN | OMN vs SMN |
| 6 | GSE93939 | Bulk RNA sequencing; laser capture microdissection | Human | Post-mortem | Healthy/non-ALS | Onuf's nucleus | Onuf's nucleus vs SMN |
| 6 | GSE93939 | Bulk RNA sequencing; laser capture microdissection | Human | Post-mortem | ALS | OMN | OMN vs SMN |

## References

1 Mead, R. J., Shan, N., Reiser, H. J., Marshall, F. & Shaw, P. J. Amyotrophic lateral sclerosis: a neurodegenerative disorder poised for successful therapeutic translation. Nat Rev Drug Discov 22, 185–212 (2023).

2 Al-Chalabi, A., van den Berg, L. H. & Veldink, J. Gene discovery in amyotrophic lateral sclerosis: implications for clinical management. Nat Rev Neurol 13, 96–104 (2017).

3 Brown, R. H. & Al-Chalabi, A. Amyotrophic Lateral Sclerosis. N Engl J Med 377, 162–172 (2017).

4 Boylan, K. Familial Amyotrophic Lateral Sclerosis. Neurol Clin 33, 807–830 (2015).

5 Fischer, L. R. et al. Amyotrophic lateral sclerosis is a distal axonopathy: evidence in mice and man. Exp Neurol 185, 232–240 (2004).

6 Comley, L. et al. Motor neurons with differential vulnerability to degeneration show distinct protein signatures in health and ALS. Neuroscience 291, 216–229 (2015).

7 Ovsepian, S. V., O’Leary, V. B. & Martinez, S. Selective vulnerability of motor neuron types and functional groups to degeneration in amyotrophic lateral sclerosis: review of the neurobiological mechanisms and functional correlates. Brain Struct Funct 229, 1–14 (2024).

8 Galtrey, C. M., Kwok, J. C., Carulli, D., Rhodes, K. E. & Fawcett, J. W. Distribution and synthesis of extracellular matrix proteoglycans, hyaluronan, link proteins and tenascin-R in the rat spinal cord. Eur J Neurosci 27, 1373–1390 (2008).

9 Jager, C. et al. Perineuronal and perisynaptic extracellular matrix in the human spinal cord. Neuroscience 238, 168–184 (2013).

10 Oohashi, T., Edamatsu, M., Bekku, Y. & Carulli, D. The hyaluronan and proteoglycan link proteins: Organizers of the brain extracellular matrix and key molecules for neuronal function and plasticity. Exp Neurol 274, 134–144 (2015).

11 Irvine, S. F. & Kwok, J. C. F. Perineuronal Nets in Spinal Motoneurones: Chondroitin Sulphate Proteoglycan around Alpha Motoneurones. Int J Mol Sci 19 (2018).

12 Morawski, M., Bruckner, G., Arendt, T. & Matthews, R. T. Aggrecan: Beyond cartilage and into the brain. Int J Biochem Cell Biol 44, 690–693 (2012).

13 Suttkus, A. et al. Aggrecan, link protein and tenascin-R are essential components of the perineuronal net to protect neurons against iron-induced oxidative stress. Cell Death Dis 5, e1119 (2014).

14 Carstens, K. E., Phillips, M. L., Pozzo-Miller, L., Weinberg, R. J. & Dudek, S. M. Perineuronal Nets Suppress Plasticity of Excitatory Synapses on CA2 Pyramidal Neurons. J Neurosci 36, 6312–6320 (2016).

15 Fawcett, J. W., Oohashi, T. & Pizzorusso, T. The roles of perineuronal nets and the perinodal extracellular matrix in neuronal function. Nat Rev Neurosci 20, 451–465 (2019).

16 Reichelt, A. C., Hare, D. J., Bussey, T. J. & Saksida, L. M. Perineuronal Nets: Plasticity, Protection, and Therapeutic Potential. Trends Neurosci 42, 458–470 (2019).

17 Balmer, T. S. Perineuronal Nets Enhance the Excitability of Fast-Spiking Neurons. eNeuro 3 (2016).

18 Wen, T. H., Binder, D. K., Ethell, I. M. & Razak, K. A. The Perineuronal ‘Safety’ Net? Perineuronal Net Abnormalities in Neurological Disorders. Front Mol Neurosci 11, 270 (2018).

19 Gray, E., Thomas, T. L., Betmouni, S., Scolding, N. & Love, S. Elevated matrix metalloproteinase-9 and degradation of perineuronal nets in cerebrocortical multiple sclerosis plaques. J Neuropathol Exp Neurol 67, 888–899 (2008).

20 Bouisset, G., Tixier, F. J., Dupak, T., Lejards, C. & Verret, L. Enriched environment requires remodeling of hippocampal perineuronal nets to trigger memory improvement in Alzheimer’s mouse model. iScience 113344 (2025).

21 Cheung, S. W., Bhavnani, E., Simmons, D. G., Bellingham, M. C. & Noakes, P. G. Perineuronal nets are phagocytosed by MMP-9 expressing microglia and astrocytes in the SOD1(G93A) ALS mouse model. Neuropathol Appl Neurobiol 50, e12982 (2024).

22 Cheung, S. W., Willis, E. F., Simmons, D. G., Bellingham, M. C. & Noakes, P. G. Phagocytosis of aggrecan-positive perineuronal nets surrounding motor neurons by reactive microglia expressing MMP-9 in TDP-43(Q331K) ALS model mice. Neurobiol Dis 200, 106614 (2024).

23 Mizuno, H., Warita, H., Aoki, M. & Itoyama, Y. Accumulation of chondroitin sulfate proteoglycans in the microenvironment of spinal motor neurons in amyotrophic lateral sclerosis transgenic rats. J Neurosci Res 86, 2512–2523 (2008).

24 Arranz, A. M. et al. Hyaluronan deficiency due to Has3 knock-out causes altered neuronal activity and seizures via reduction in brain extracellular space. J Neurosci 34, 6164–6176 (2014).

25 Saghatelyan, A. K. et al. Reduced perisomatic inhibition, increased excitatory transmission, and impaired long-term potentiation in mice deficient for the extracellular matrix glycoprotein tenascin-R. Mol Cell Neurosci 17, 226–240 (2001).

26 Freitag, S., Schachner, M. & Morellini, F. Behavioral alterations in mice deficient for the extracellular matrix glycoprotein tenascin-R. Behav Brain Res 145, 189–207 (2003).

27 Fukamauchi, F. et al. Abnormal behavior and neurotransmissions of tenascin gene knockout mouse. Biochem Biophys Res Commun 221, 151–156 (1996).

28 Kaplan, A. et al. Neuronal matrix metalloproteinase-9 is a determinant of selective neurodegeneration. Neuron 81, 333–348 (2014).

29 Collins, M. A., An, J., Hood, B. L., Conrads, T. P. & Bowser, R. P. Label-Free LC-MS/MS Proteomic Analysis of Cerebrospinal Fluid Identifies Protein/Pathway Alterations and Candidate Biomarkers for Amyotrophic Lateral Sclerosis. J Proteome Res 14, 4486–4501 (2015).

30 Dityatev, A. et al. Activity-dependent formation and functions of chondroitin sulfate-rich extracellular matrix of perineuronal nets. Dev Neurobiol 67, 570–588 (2007).

31 Geissler, M. et al. Primary hippocampal neurons, which lack four crucial extracellular matrix molecules, display abnormalities of synaptic structure and function and severe deficits in perineuronal net formation. J Neurosci 33, 7742–7755 (2013).

32 Giamanco, K. A. & Matthews, R. T. Deconstructing the perineuronal net: cellular contributions and molecular composition of the neuronal extracellular matrix. Neuroscience 218, 367–384 (2012).

33 Dickens, S., Goodenough, A. & Kwok, J. An in vitro neuronal model replicating the in vivo maturation and heterogeneity of perineuronal nets. BioRxiv (2022).

34 Maury, Y. et al. Combinatorial analysis of developmental cues efficiently converts human pluripotent stem cells into multiple neuronal subtypes. Nat Biotechnol 33, 89–96 (2015).

35 Bryson, J. B. et al. Optical control of muscle function by transplantation of stem cell-derived motor neurons in mice. Science 344, 94–97 (2014).

36 Harley, P. et al. Aberrant axon initial segment plasticity and intrinsic excitability of ALS hiPSC motor neurons. Cell Rep 42, 113509 (2023).

37 Machado, C. B. et al. In Vitro Modelling of Nerve-Muscle Connectivity in a Compartmentalised Tissue Culture Device. Adv Biosyst 3 (2019).

38 Kwok, J. C., Carulli, D. & Fawcett, J. W. In vitro modeling of perineuronal nets: hyaluronan synthase and link protein are necessary for their formation and integrity. J Neurochem 114, 1447–1459 (2010).

39 Jakovljevic, A. et al. Structural and Functional Modulation of Perineuronal Nets: In Search of Important Players with Highlight on Tenascins. Cells 10 (2021).

40 Ziff, O. J. et al. Integrated transcriptome landscape of ALS identifies genome instability linked to TDP-43 pathology. Nat Commun 14, 2176 (2023).

41 Ziff, O. J. et al. Meta-analysis of human and mouse ALS astrocytes reveals multi-omic signatures of inflammatory reactive states. Genome Res 32, 71–84 (2022).

42 Lemieux, S. P., Lev-Ram, V., Tsien, R. Y. & Ellisman, M. H. Perineuronal nets and the neuronal extracellular matrix can be imaged by genetically encoded labeling of HAPLN1 in vitro and in vivo. bioRxiv (2023).

43 Wegrzyn, D., Freund, N., Faissner, A. & Juckel, G. Poly I:C Activated Microglia Disrupt Perineuronal Nets and Modulate Synaptic Balance in Primary Hippocampal Neurons in vitro. Front Synaptic Neurosci 13, 637549 (2021).

44 Tewari, B. P., Chaunsali, L., Prim, C. E. & Sontheimer, H. A glial perspective on the extracellular matrix and perineuronal net remodeling in the central nervous system. Front Cell Neurosci 16, 1022754 (2022).

45 Asher, R. A. et al. Versican is upregulated in CNS injury and is a product of oligodendrocyte lineage cells. J Neurosci 22, 2225–2236 (2002).

46 Mehta, A. R. et al. Mitochondrial bioenergetic deficits in C9orf72 amyotrophic lateral sclerosis motor neurons cause dysfunctional axonal homeostasis. Acta Neuropathol 141, 257–279 (2021).

47 Barton, S. K. et al. Transactive response DNA-binding protein-43 proteinopathy in oligodendrocytes revealed using an induced pluripotent stem cell model. Brain Commun 3, fcab255 (2021).

48 Fernandopulle, M. S. et al. Transcription Factor-Mediated Differentiation of Human iPSCs into Neurons. Curr Protoc Cell Biol 79, e51 (2018).

