## Supplementary code 1 for "ALS mutations do not alter perineuronal net formation in human stem cell-derived motor neurons"

Macro script written for use in FIJI software to threshold images of PNN staining. Script was applied to original images obtained using a Zeiss LSM 980 confocal microscope.

```
// ask user to select a folder

dir1 = getDirectory("C:/Users/Kerin/Downloads");

// get the list of files (& folders) in it

list = getFileList(dir1);

// prepare a folder to output the images

dir2= dir1 + File.separator + "Overlay and PNN stain" + File.separator ;

File.makeDirectory(dir2);

//setBatchMode(true);

for (i=0; i<list.length; i++) {

showProgress(i+1, list.length);

open(dir1+list[i]);

run("Z Project...", "projection=[Max Intensity]");

title = getTitle();

run("Split Channels");

// Run threshold and apply colour to each z stack image for each channel

title1 = "C1-" + title;

selectWindow(title1); {

setAutoThreshold("Default dark");

//run("Threshold...");

setThreshold(47, 255);

setOption("BlackBackground", false);

run("Convert to Mask");

//run("Channels Tool...");

run("Cyan");

//run("Flatten");
```

```
// rename("TUJ1");
//TUJ1 = getTitle();
// saveAs("TIFF", output_dir_composite + TUJ1);
}
title2 = "C2-" + title;
selectWindow(title2); {
setAutoThreshold("Default dark");
//run("Threshold...");
setThreshold(50, 255);
setOption("BlackBackground", false);
run("Convert to Mask");
//run("Channels Tool...");
run("Green");
// run("Flatten");
//rename("DAPI");
//DAPI = getTitle();
// saveAs("TIFF", output_dir_composite + DAPI);
}
title3 = "C3-" + title;
selectWindow(title3); {
setAutoThreshold("Default dark");
//run("Threshold...");
setThreshold(50, 255);
setOption("BlackBackground", false);
run("Convert to Mask");
//run("Channels Tool...");
run("Red");
//rename("Aggrecan or other PNN staining name");
```

```
//Aggrecan = getTitle();

//saveAs("TIFF", dir2 + title);

}

// Run overlay

//Here overlay DAPI on TUj1 first and then add ACAN over DAPI/TUj image

selectWindow(title3);

saveAs("TIFF", dir2 + title3);

selectWindow(title1);

run("Add Image...", "image=[&title2] x=0 y=0 opacity=50 zero");

rename("TUJ_DAPI" + title1);

run("Flatten");

saveAs("TIFF", dir2 + "TUJ_DAPI" + title1);

run("Add Image...", "image=[&title3] x=0 y=0 opacity=50 zero");

rename("W_PNN_Stain" + title1);

run("Flatten");

saveAs("TIFF", dir2 + "W_PNN_Stain" + title1);

}

run("Close All");
```
