## Supplementary code 2 for "ALS mutations do not alter perineuronal net formation in human stem cell-derived motor neurons"

Macro script written for use in FIJI software to threshold images of TDP-43 staining in all cell lines. Script was applied to original images obtained using Zeiss LSM 980 confocal microscope.

```
// ask user to select a folder

dir = getDirectory("C:/Users/Kerin/Desktop/Figures/PNN counts");

// get the list of files (& folders) in it

fileList = getFileList(dir);

// prepare a folder to output the images

output_dir_splits = dir + File.separator + " output_splits_Threshold" + File.separator ;

File.makeDirectory(output_dir_splits);

//activate batch mode

setBatchMode(true);

// LOOP to process the list of files

for (i = 0; i < lengthOf(fileList); i++) {

// define the "path"

// by concatenation of dir and the i element of the array fileList

current_imagePath = dir+fileList[i];

// check that the currentFile is not a directory

if (!File.isDirectory(current_imagePath)){

// open the image and split

open(current_imagePath);

// get some info about the image

getDimensions(width, height, channels, slices, frames);

//setAutoThreshold("Default");

run("Threshold...");

setThreshold(0, 30);

setOption("BlackBackground", false);

run("Convert to Mask");
```

```
title = getTitle();

saveAs("tiff", output_dir_splits +title);

}

// make sure to close every images befores opening the next one

run("Close All");

}

setBatchMode(false);
```
